# Syncytins enable novel possibilities to transduce human or mouse primary B cells and to achieve well-tolerated *in vivo* gene transfer

**DOI:** 10.1101/816223

**Authors:** Y. Coquin, M. Ferrand, A. Seye, L. Menu, A. Galy

## Abstract

Syncytins are cellular transmembrane glycoproteins with fusogenic and immunosuppressive properties that are encoded by endogenous retroviral envelope sequences in mammalian genomes. Based on their properties, syncytins may be useful to pseudotype lentiviral gene transfer vectors (LV) and to obtain well-tolerated *in vivo* gene delivery but their cellular targets are unknown in this context. We pseudotyped LV with human or murine syncytins. Such LV-Syn particles were infectious *in vitro* but required a transduction additive, as do other retroviral envelope LV pseudotypes. In these conditions, LV-Syn remarkably transduced quiescent human or murine primary B cells at high level *in vitro* including naïve blood B cells or B cell precursors from murine bone marrow. Transduced human B cells could be expanded in culture and were functional. Human or murine T cells were transduced less efficiently than B cells, in agreement with lower levels of syncytin receptors on T cells compared to B cells. Well-tolerated *in vivo* gene transfer was possible without additive, as demonstrated with murine syncytin A-mediated gene delivery in C57BL/6 mice. A single intravenous injection of LV-SynA vector to mice led to stable gene transfer into spleen germinal center B cells. LV-SynA were also intrinsically less immunogenic than LV-VSVG, leading to low antibody responses against the vector capsid. This is the first evidence of interactions between syncytins and B cells, providing novel opportunities for B cell genetic engineering and for well-tolerated gene transfer *in vivo*. The findings also suggest that some immunosuppressive properties of syncytins could be mediated by B cells.

**One Sentence Summary:** Syncytins are fusogenic cellular proteins that can pseudotype lentiviral gene transfer vector particles, achieving efficient gene transfer into primary quiescent B cells and reducing the *in vivo* immunogenicity of the particles following systemic administration.

## Introduction

Efficient viral vectors for gene transfer have enabled the recent clinical successes of gene therapy in genetic diseases or cancer. Yet, the *in vivo* use of viral vectors can be limited by adaptive immune recognition of their components. Advanced self-inactivated HIV-1-derived lentiviral vectors (LV) are powerful vectors that stably transduce quiescent cells, integrate safely into the host cell chromatin and permit long-term engraftment of transduced cells (reviewed in (1–3). The entry of LV into target cells is mediated by envelope glycoproteins which interact with cellular receptors on target cells and induce cell:virus membrane fusion. The most commonly-used envelope glycoprotein to pseudotype LV is the vesicular stomatitis G glycoprotein (VSVG) enabling gene delivery into a broad range of target cells with exception of certain cell types such as quiescent human T or B cells which poorly express the cognate LDL receptors (4). LV are largely employed for *ex vivo* gene therapy and *in vivo* they are used locally in the brain or eye where immune reactions are limited. However, VSVG is immunogenic and the systemic administration of LV-VSVG to C57BL6 mice leads to the transduction of macrophages in the liver and of dendritic cells in lymphoid tissues, inducing inflammation and strong CD4 and CD8 T cell immune responses against transgene products (5–7). Envelope glycoproteins of LV are key determinants of their interactions with the immune system. VSVG can be shielded with CD47 to prevent uptake by liver Kupfer cells and to reduce inflammatory signals *in vivo* (7). Alternatively, various LV pseudotypes exist, many of which have been elaborated from mutated and engineered viral envelope glycoproteins to increase cell-specific targeting in particular towards B cells or to permit *in vivo* LV delivery, as reviewed recently (8). Yet these systems which are always based on the use of viral platforms are not devoid of immunogenic potential. Possibly, replacing the exposed viral components of LV envelopes by endogenous cellular components to which the host is tolerant could reduce immunogenicity but this has not been effectively obtained or tested in immunocompetent hosts.

During evolution, endogenous retroviruses (ERV) have entered the germline of mammalian hosts. Some ERV genes have become endogenized, persisting over tens of millions of years in mammalian genomes as they confer physiological functions (9). This is the case of syncytin genes which are cellular genes corresponding to functional ERV env open reading frames found as single copies, and which promote the formation of the syncytiotrophoblast during placentation (10–13). Structurally, syncytins resemble typical retroviral env glycoproteins produced by cleavage of a precursor polypeptide into two mature proteins that associate as homotrimers. A surface subunit serves for receptor recognition and a transmembrane subunit bears a fusion peptide and an immuno-suppressive domain (14). Various syncytin genes have been identified in the genomes of all mammalian species examined, resulting from independent gene captures (13). In the human genome, two syncytin genes ERVW-1 (syncytin 1 or Syn1) and ERVFRD-1 (syncytin 2 or Syn2) have entered the primate lineage between 25-40 million years ago. They interact respectively with membrane receptors identified as the neutral amino acid transporter ASCT-2 (SLC1A5 gene) and ASCT-1 (SLC1A4 gene) for Syn1, and the phospholipid transporter/symporter MFSD-2A for Syn-2 (15, 16) (17). In mice, syncytin-A (SynA) and syncytin-B (SynB) which are phylogenetically unrelated to human Syn1 and Syn2 have entered the Muridae lineage more than 20 million years ago. Murine syncytins are also fusogenic, highly expressed in placenta (11) and required for its development, SynA being critically-required to maintain gestation (18). Recently, Ly6e, a GPI-anchored membrane protein was identified as the receptor for SynA (19) while the receptor for SynB remains unknown. Outside of the placenta, syncytins also contribute to other important membrane fusion processes as they are up-regulated in regenerating skeletal muscle after strong exercise (20) and SynB promotes myoblast fusion and muscle formation, contributing to the male muscle sexual dimorphism in mice (21). SynB also contributes to the formation of multinucleated osteoclasts without being essential for bone homeostasis (22). Some syncytins are endowed with immunosuppressive properties which are linked to immuno-suppressive domain motifs (23) but these immunosuppressive mechanisms are not well known and not as conserved as the fusogenic capacity of syncytins.

Syncytins have not yet been practically employed to pseudotype gene transfer vectors. Syn1-pseudotyped recombinant HIV-derived LV have been reported, but titers were low and further reduced by concentration, freezing and thawing rendering the preparations useless (24). A deletion of the R intracytoplasmic region increased the fusion efficiency and infectious titers but this was not practically useful (15). Syn2 could pseudotype SIV, HIV or MLV vectors but very low titers also precluded functional studies (25). Yet, syncytins remain attractive cellular genes for LV pseudotype as they may facilitate *in vivo* use. This prompted us to investigate the potential of human Syn1 and Syn2 as well as murine SynA and SynB to pseudotype LV. To optimize transduction, we used Vectofusin-1 (VF1), a peptidic transduction additive identified in our laboratory. VF1 is a short histidine-rich amphipathic peptide known to enhance the infectivity of lentiviral or gamma-retroviral vectors pseudotyped with various retroviral envelope glycoproteins, for instance Gibbon ape leukemia virus, the feline endogenous virus RD114 or baboon endogenous virus (BaEV) glycoproteins (26–28). VF1 enhances adhesion of LV to target cells, aggregates and pulls-down particules towards cells and promotes virus:cell membrane fusion. Here, the use of VF1 significantly enhanced syncytin-mediated transduction revealing the cellular tropism of syncytins for human or murine primary B lymphocytes. LV pseudotyped with SynA could also be used *in vivo* and induced less antibody immune responses than LV-VSVG.

## Results

### Syncytin-pseudotyped LVs are infectious in the presence of VF1 and demonstrate cell-selective infectivity

To explore the use of human or murine synyctins as LV pseudotype, we transiently transfected the full-length native cDNAs of Syn1, Syn2, SynA or SynB in 293T cells to produce recombinant particles with each of these envelopes (LV-Syn1, 2, A, B). In parallel LV-VSVG wa produced to serve as a comparative control. In the conditions defined and following optimization of plasmid amounts (data not shown), it was possible to produce stable and infectious particles pseudotyped by either of the 4 syncytins (Table 1). Such LV-Syn particles were successfully concentrated by ultracentrifugation similarly to LV-VSVG (29). While LV-VSVG batches can be generated by pooling 2 consecutive harvests from the same producing cells 24 hours apart, only 1 harvest was used to produce LV-Syn because syncytin-transfected 293 T cells started to fuse after 24 hours, reducing viral titers (data not shown). The concentrated LV-Syn vectors were cryopreserved at −80°C and the particles were stable for several months (data not shown). Regardless of the syncytin used, LV-Syn had similar physical titers ranging from 4.8 to 7.2 E+05 ng p24/mL and 3.5 to 4.7 E+11 physical particles (pp)/mL using a particle counter (Table 1). LV-Syn physical titers were similar to those of LV-VSVG indicating that synyctins enable efficient pseudotyping in the rHIV lentiviral system.

**Table 1.** Titers of syncytin-pseudotyped LVs. Legend: Physical titers were measured by p24 ELISA and by direct physical particle (pp) counting using a particle counter. Infectious titers were measured on A20.IIa cells in the presence of VF1 and defined as infectious genomes (IG) /mL using qPCR.

| vector type | number of batches titered | ng p24/mL x E+05 | pp/mL x E+11 | IG/mL x E+06 |
| --- | --- | --- | --- | --- |
| LV-Syn1 | *n*=3 | 7.2 ± 2.2 | 4.0 ± 2.7 | 4.4 ± 2.0 |
| LV-Syn2 | *n*=5 | 5 ± 4.0 | 4.5 ± 2.6 | 5.0 ± 2.8 |
| LV-SynA | *n*=12 | 10 ± 1.3 | 4.7 ± 4.6 | 11 ± 0.8 |
| LV-SynB | *n*=4 | 4.8 ± 5.5 | 3.5 ± 3.5 | 10 ± 0.9 |
| LV-VSVg | *n*=3 | 2.1 ± 2.0 | 2.1 ± 1.5 | 22 ± 2.5 |

Infectious properties of LV-Syn were demonstrated with several transgenes including GFP or truncated NGFR and with the use of the VF1 transduction additive (Figure 1 A-C). Initially we tested the human BeWo choriocarcinoma cell line known to express both the ASCT2 and MFSD2a receptors of Syn1 and Syn2 (17). Little transduction was obtained by vectors used alone or in the presence of cationic agents such as protamine sulfate (PS) or polybrene (PB), two agents that are commonly used to facilitate LV cellular entry (Supplementary Figure S1A). However, BeWo cell transduction was markedly enhanced by adding VF1 (Supplementary Figure S1A), prompting a systematic testing of this peptide on various cell lines. In the presence of VF1, human 293T cells were transduced efficiently by human LV-Syn vectors (Figure 1A) resulting in vector dose-dependent transgene expression, stable expression over time well-correlated to proviral integration (Supplementary Figures S1 B-E). Murine A20.IIa lymphoma cells were transduced very efficiently by murine syncytins in the presence of VF1 (Figure 1B). However, VF1 did not have a universal effect. For instance, murine syncytin-pseudotyped LVs did not transduce 293T cells even in the presence of VF1 (Figure 1C), therefore only human syncytin-pseudotyped LVs could be titered on 293T cells. The effects of syncytins were not species-restricted (Figure 1C). Cells lines such as 293T cells or Jurkat T cells were transduced by some but not all of the LV-Syn, while others could not be transduced by any of the human or murine LV-Syn tested. Refractory cells included human HCT116 cells which are routinely used in our laboratory for the titration of LV-VSVG (29), murine C2C12 myoblasts or NIH3T3 murine fibroblasts. On the contrary, A20.IIa murine lymphoma cells were readily transduced by all of the LV-Syn tested (Figure 1C). Raji B cells were also transduced although not as efficiently as A20.IIa cells, but FACS-sorted transgene-positive Raji cells continued to stably express the transgene and displayed vector integration for at least 49 days of continuous *in vitro* culture (data not shown). Since A20.IIa cells enabled the highest levels of infection by all LV-Syn tested and were also permissive to LV-VSVg, these cells were chosen to establish an infectious genome (IG) titration by qPCR. When compared, these titers in IG/mL on A20.a cells were similar as those done as TU/mL by FACS on 293T cells (data not shown). Human or murine syncytin-pseudotyped LV were found to have comparable infectious titers, ranging from 4.4 to 11 E+06 IG/mL (Table 1). In the conditions used for production, LV-VSVG batches titered 2 to 5 times higher than LV-Syn batches. Thus, LV can be efficiently pseudotyped with human or murine syncytins and the particles are functionally infectious in the presence of an adjuvant.

**Fig. 1:**
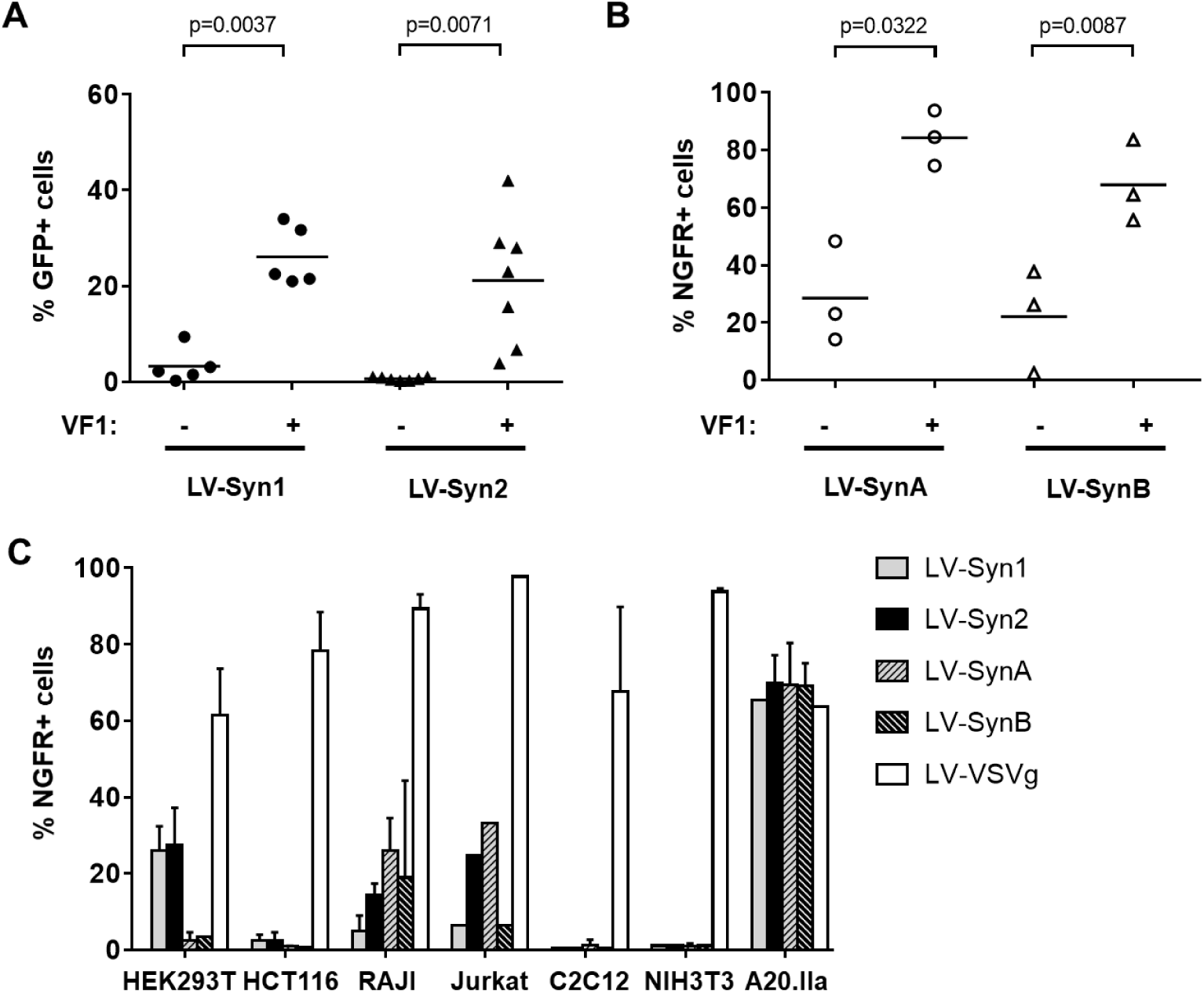
Gene transfer into various cell lines with syncytin-pseudotyped vectors and VF1. (**A**). Human 293T cells were infected with 1E+05 TU/mL of LV-Syn1 or LV-Syn2 encoding GFP in the presence or not of VF1. GFP expression was measured after 3 days by FACS. The enhancing effect of VF1 was statistically significant as shown in 5-7 experiments using a paired t test. (**B**). Murine A20.IIa cells were infected with 1E+05 IG/mL LV-SynA or LV-SynB encoding a truncated NGFR, in the presence or not of VF1. NGFR expression was measured after 3 days by FACS. The enhancing effect of VF1 was statistically significant as shown in 3 experiments using a paired t test. (**C**). Various human (HEK293T, HCT116, RAJI, Jurkat) or murine (C2C12, NIH3T3, A20.IIa) cell lines were infected with 1E+05 IG/mL of the indicated vectors encoding NGFR in the presence of VF1. NGFR was measured after 7 days by FACS. Results are representative of 3 separate experiments.

### LV-Syn1 and LV-Syn2 vectors efficiently transduce human peripheral blood B cells

Positive results on lymphoid cell lines led us to evaluate the transduction of primary blood lymphocytes by LV-Syn. Human peripheral blood mononuclear cell (PBMC) were infected by LV-Syn1 or LV-Syn2 vectors either immediately after cellular isolation, or following a short culture in the presence of IL-7 aiming to maintain the viability of naive lymphocytes overnight (schema in Supplementary Figure S2A). The total CD45+ nucleated cell population showed transgene expression when VF1 was added and IL-7 pre-activation increased these levels to about 13% and 5% respectively with LV-Syn1 or LV-Syn2 (Supplementary Table S1). However, among these cells, the small subsets of CD19+ B lymphocytes and of CD11c+ cells expressed high levels of transgene whereas only a fraction of the more abundant CD3+ T cells were marked (Figure 2-top panels, and Table 2). An average of 59% and 44% of CD19+ cells were transduced by LV-Syn1 or LV-Syn2 respectively following IL-7 pre-activation and in the presence of VF1. Such levels were at least as high or even higher than those obtained with high concentrations of LV-VSVG (Table 2). In contrast, peripheral blood T cell marking were not consistent from experiment to experiment but were low (Figure 2 and Table 2). Without prior activation, almost no transduction of T cells is observed.

**Table 2:** Transduction of human peripheral blood lymphocytes with syncytin-pseudotyped LV. Legend: Human PBMC were transduced with the indicated vectors and with or without VF1 in steady-state conditions (no activation) or following a short overnight pre-activation with IL-7, and subsequently cultured with IL-7 for 3 days. LV-Syn vectors were used at 1E+05 TU/mL after FACS titration on 293T cells and LV-VSVg were used at 1E+08IG/mL as titered by qPCR on HCT116 cells. Transduction results are expressed as the average percentage ± SD of cells positive for GFP in CD19+ (B cells), CD3+ (T cells) or CD11c+ (myeloid dendritic cells). Data represent 3 separate experiments without VF1 and 8 to 9 separate experiments in the presence of VF1. (*) p<0.05 compared to “LV-VSVG+VF1” in a paired Student’s t test

| % GFP+ cells | no activation |  |  | IL-7 activation |  |  |
| --- | --- | --- | --- | --- | --- | --- |
|  | CD19+<br>B cells | CD3+<br>T cells | CD11c+<br>Myeloid<br>cells | CD19+<br>B cells | CD3+<br>T cells | CD11c+<br>Myeloid<br>cells |
| PBS | 0.0±0 | 0.1±0.1 | 0.2±0.3 | 0.3±0.4 | 0.5±0.4 | 0.8±0.6 |
| LV-Syn1 | 2.3±4 | 1.3±2.6 | 5.5±6.3 | 2.1±2.6 | 0.8±0.3 | 4.1±6.0 |
| LV-Syn2 | 0.6±0 | 0.4±1.0 | 0.2±8.2 | 1.8±1.5 | 1.5±1.0 | 2.2±2.2 |
| LV-VSVg | 3.6±5 | 2.8±2.9 | 11.5±3.5 | 2.6±1.6 | 1.3±0.6 | 2.4±0.8 |
| VF1 | 3.3±4 | 0.24±0.2 | 7.8±8.4 | 0.8±0.5 | 0.18±0.2 | 0.25±0.5 |
| LV-Syn1 + VF1 | 35.7± | 3.61±3.6 | 57.7±11 | 59.3±23 * | 11.2±19 | 59.2±18 |
| LV-Syn2 + VF1 | 22.2± | 1.49±1.9 | 35.5±34 | 44.0±24 | 6.1±10 | 40.1±15 |
| LV-VSVg + VF1 | 27.6± | 0.97±0.6 | 45.2±16 | 32.1±18 | 3.5±1.4 | 35.1±10 |

**Fig. 2:**
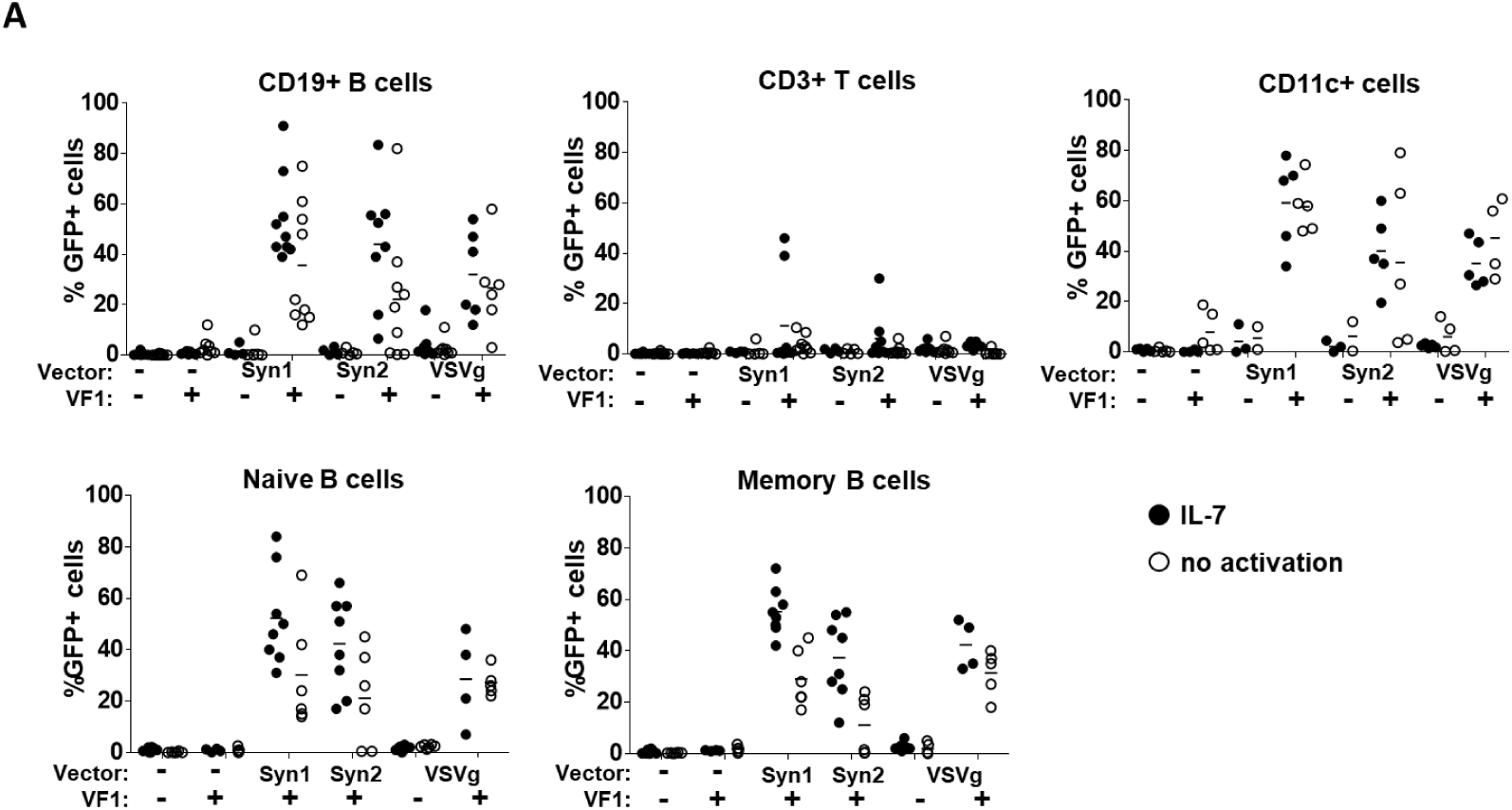
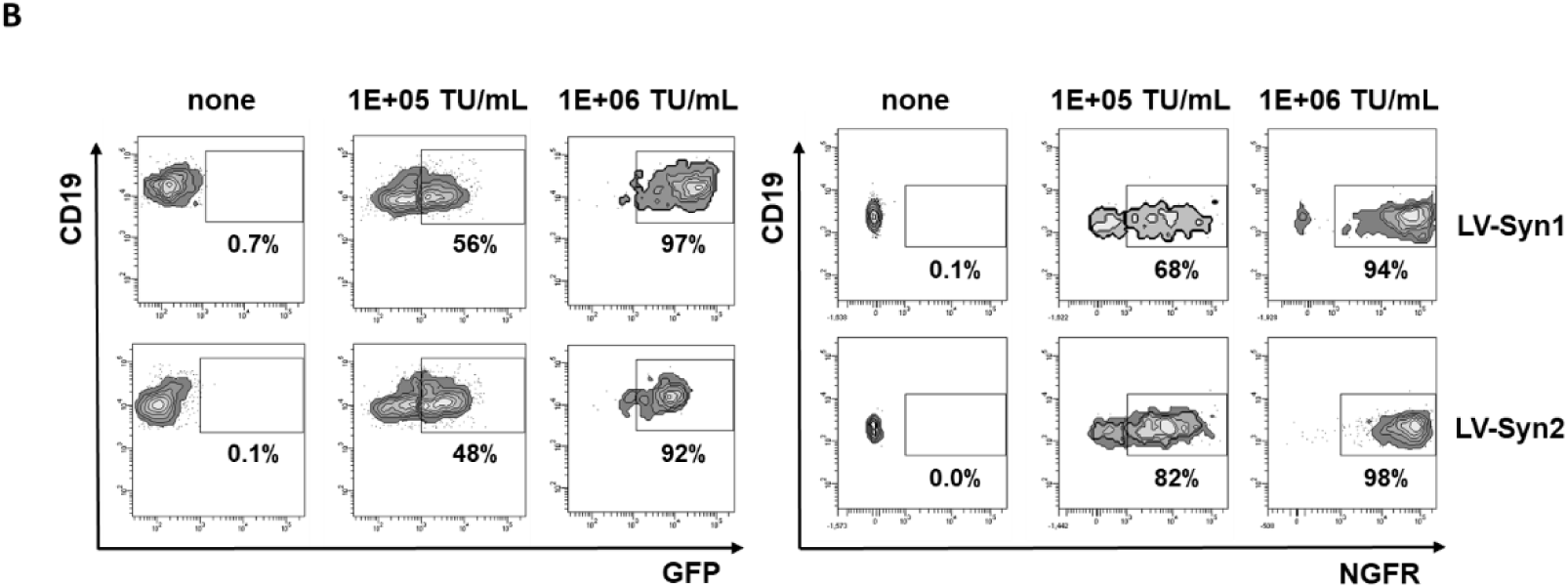
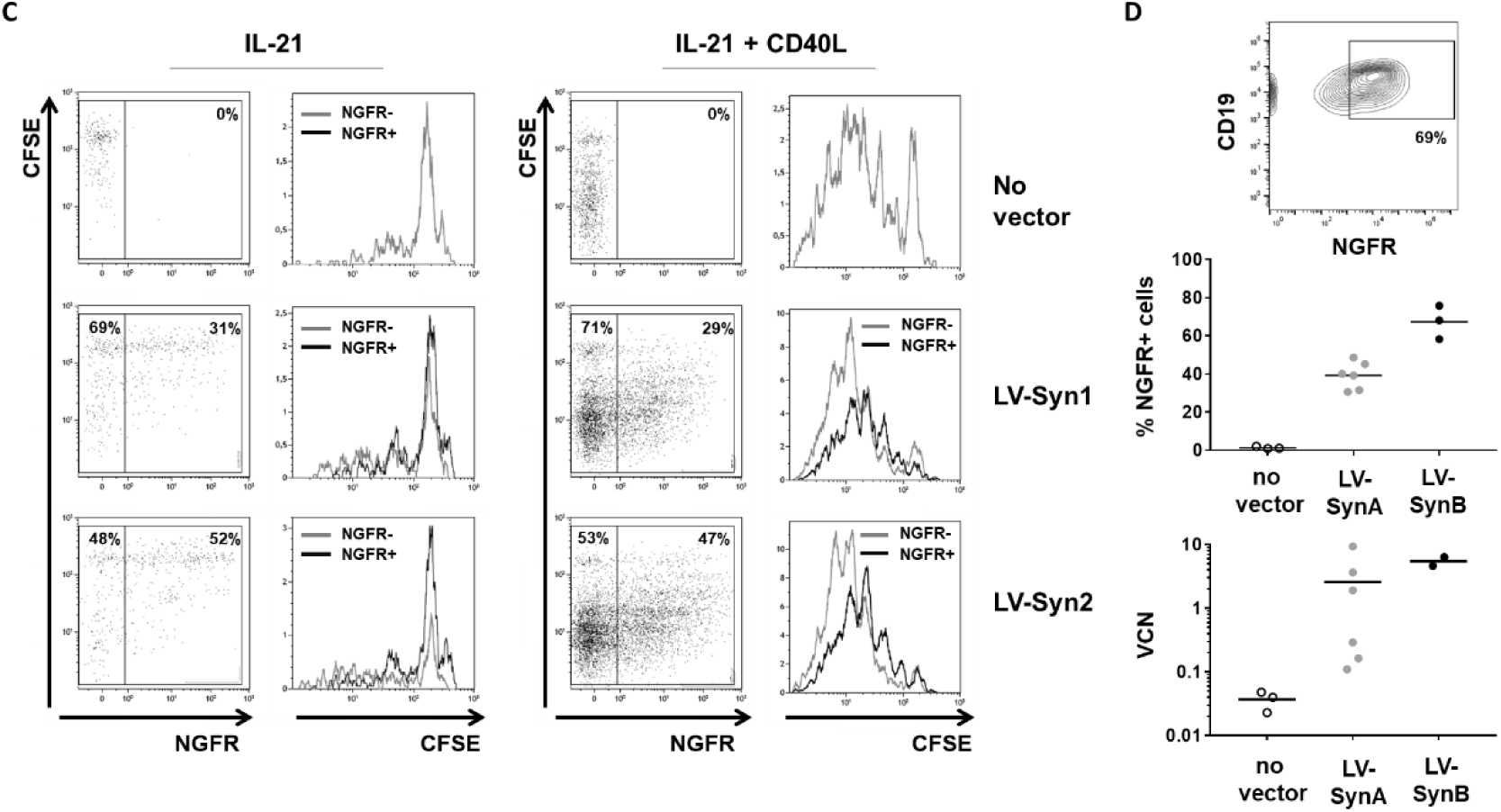
Transduction of primary blood cell subsets with syncytin-pseudotyped vectors. (**A**). Human peripheral blood mononuclear cells (PBMC) (4 to 9 separate blood donors) were infected with GFP-encoding LV-Syn1 or LV-Syn2 vectors (1E+05 TU/mL) in the absence or presence of VF1 and either immediately after isolation of the cells (no activation – open symbols) or following an overnight culture of cells with IL-7 (10 ng/mL)(closed symbols). After 3 days, GFP was measured by FACS in different subsets of live cells (CD3+ CD19-CD11c-T cells; CD3-CD19+ CD11c-B cells; CD3-CD19-CD11c+ myeloid dendritic cells), naive (CD27-) or memory (CD27+) B cell subsets. (**B**). Representative results of 3 PBMC donors incubated with IL-7 overnight, then transduced with NGFR-encoding LV-Syn1 or LV-Syn2 at concentrations 1E+05 and 1E+06 TU/mL in the presence of VF1. Transgene expression was measured by FACS after 3 days. (**C**). PBMCs were labeled with 2µM CFSE, transduced with LV-Syn1 or LV-Syn2 vectors encoding NGFR, and cultured in the absence or presence of CD40 Ligand (CD40L) (2µg/mL) and/or IL-21 (50ng/mL). After 3 days culture, B cell transduction (expression of NGFR on CD19+ cells) and B cell proliferation (CFSE dilution) were determined by flow cytometry. Results representative of 6 PBMC donors tested in 2 separate experiments (3 PBMC donors each). (**D**). CD19+ cells were sorted from PBMC (3 donors) and were transduced with LV-SynA or LV-SynB encoding NGFR (1E+05 IG/mL) and transgene expression was analyzed by FACS after 7 days.

Since it was possible to transduce quiescent non-activated CD19+ cells, further investigation of CD19+ cell subsets (30) was undertaken using a FACS gating strategy shown in Supplementary Figure S2B. Both naive and memory B cell subsets were effectively transduced with the human syncytin-pseudotyped vectors in the presence of VF1, regardless of prior activation (Figure 2 – bottom panels). Naive CD19+ CD27-IgD+ cells (representing 50-70% of the CD19+ B cell population); memory non-switched CD19+ CD27+ IgD+ cells (15-20% of the population); memory switched CD19+ CD27+ IgD-IgM-cells (2-5% of the population); memory switched IgM only CD19+ CD27+ IgD-IgM+ cells (2-5% of the population) and plasmablasts CD19+ CD27hi (1-2% of the population) were transduced efficiently in the absence of prior activation and more effectively than LV-VSVG suggesting a specific tropism of the syncytins for quiescent B cells (Table 3). The transduction of CD19+ B cells by LV-Syn1 and LV-Syn2 were vector concentration-dependent and transgene-independent (Figure 2B). High concentrations of vector enabled the transduction of almost all peripheral blood CD19+ cells to express either GFP or NGFR, thereby providing an effective system to express heterologous proteins in these cells.

**Table 3:** Percentage of transduction of B cells subsets. Legend: Human PBMC were transduced in steady-state conditions with LV-Syn1 or LV-Syn2 (1E+05 TU/mL) or LV-VSVg (1E+07 TU/mL). Results are the average percentage ± SD of cells positive for GFP in the indicated populations defined as naive B cells (CD3-, CD19+, CD27-, IgM+, IgD+ cells; memory non-switch B cells (CD3-, CD19+, CD27+, IgM+, IgD+ cells); memory IgM only B cells (CD3-, CD19+, CD27+, IgM+, IgD- cells); memory switched B cells (CD3-, CD19+, CD27+, IgM-, IgD-) and plasmablasts (CD3-, CD19+, CD27high), 3 days after infection. Data represent 2 to 3 separate experiments with 1 to 2 different donor per experiment. nd=not determined because too few cells to analyze in the gate.

|  | Naive | Memory non-switch | Memory IgM only | Memory switched | Plasmablasts |
| --- | --- | --- | --- | --- | --- |
| PBS | 0.2±0.3 | 0.0±0 | 0.3±0.5 | 0.5±1 | 0.3±0.6 |
| **VF1** | **0.6±0.5** | **1±0.1** | **0±0** | **0.3±0.5** | **0.3±0.6** |
| LV-Syn1 | 0.4±1.1 | 1±0.9 | 0.2±0.3 | 0±0 | 0.3±0.6 |
| **LV-Syn1+VF1** | **62±26** | **23±3.3** | **16±13** | **41±11** | **32±13** |
| LV-Syn2 | 1±0.1 | 4±5.2 | 5±6 | 4±5.3 | 1.00±1.4 |
| **LV-Syn2+VF1** | **52±29** | **29±15** | **27±34** | **26±30** | **29±8.5** |
| LV-VSVg | 0.1±0.1 | 9±8.4 | 4±4 | 10±16 | 2±1.1 |
| **LV-VSVg+VF1** | **14±14** | **18±16** | **12±11** | **20±11** | **nd** |

Transduction with syncytin-pseudotyped vectors did not appear to affect B cell division and activation. Resting human B cells transduced or not with LV-Syn1 or LV-Syn2 proliferated moderately in response to IL-21 alone and more extensively in response to IL21+CD40L as expected (31). Transgene-expressing CD19+ cells proliferated as much as the non-transduced cells (Figure 2C) (Table 4). Transduced cells were able to mature into plasmablasts antibody secreting cells expressing high levels of CD38, CD27 and membrane IgG (data not shown). The proportion of plasmablasts was similar in transduced and non-transduced B cells indicating that exposure to LV-Syn vectors did not prevent cellular activation towards terminal differentiation (Table 4).

**Table 4.** Functional B cells are stably transduced by syncytin-pseudotyped LV. Legend: Data were obtained from human PBMCs of 6 different healthy donors. Cellular division was measured by CFSE labeling following transduction with LV-Syn1 or LV-Syn2 encoding delta-NGFR. Cells were cultured in the absence or presence of CD40L [2μg/mL] and/or IL-21 [50ng/mL]. After 3 and 6 days culture, B cell transduction (ie, NGFR expression), B cell proliferation (ie, CFSE dilution) and the frequency of CD19+ CD27++ CD38+ plasmablasts were determined by FACS.

|  | Day 3 |  |  |  | Day 6 |  |  |  |
| --- | --- | --- | --- | --- | --- | --- | --- | --- |
| Percentage | IL-21 |  | IL-21 + CD40L |  | IL-21 |  | IL-21 + CD40L |  |
|  | LV-Syn1 | LV-Syn2 | LV-Syn1 | LV-Syn2 | LV-Syn1 | LV-Syn2 | LV-Syn1 | LV-Syn2 |
| **NGFR+ B cells** | 37.2 ± 5.3 | 60.2 ± 8 | 26.8 ± 9.4 | 45.8 ± 11.5 | 39.8 ± 9.4 | 45.3 ± 6.9 | 17.3 ± 2.7 | 23.3 ± 6.8 |
| **NGFR+ dividing B cells** | 25.2 ± 15.7 | 16.7 ± 6 | 77.5 ± 16.4 | 82.3 ± 10 | 28.8 ± 12.8 | 28.7 ± 14.1 | 93.2 ± 3.4 | 93.5 ± 4 |
| **NGFR-neg dividing B cells** | 35.3 ± 16.7 | 34.2 ± 17 | 85.8 ± 7.4 | 91.0 ± 4.4 | 40.5 ± 18.9 | 31.7 ± 8.6 | 94.8 ± 1.2 | 95.2 ± 1.5 |
| **plasmablasts in NGFR+ B cells** | 4.4 ± 2.0 | 5.7 ± 2.7 | 13.6 ± 3.7 | 14.1 ± 4.4 | 17.8 ± 7.9 | 18.2 ± 10.2 | 29.8 ± 13.1 | 28.9 ± 9.6 |
| **plasmablasts in NGFR-neg B cells** | 4.7 ± 1.2 | 6.0 ± 3.0 | 15.8 ± 5.7 | 14.5 ± 5.6 | 20.8 ± 14.1 | 16.3 ± 12 | 32.7 ± 15 | 30.2 ± 10.9 |

Human CD19+ B cells could also be transduced with either of the murine syncytin-pseudotyped LV, proliferated robustly in response to IL21+CD40L for 7days and expressed the transgene (Figure 2D). Both of the murine syncytins could mediate the transduction of human B cells, but based on transgene levels and integrated vector copies, syncytin B seemed to be more efficient than syncytin A, although additional experiments are needed for confirmation.

### Syncytins fail to mediate transduction of CD34+ cells

Hematopoietic stem and progenitor CD34+ cells express ASCT2, a receptor for the RD114 feline endogenous virus or for the baboon endogenous virus BaEV, and also express ASCT1 which is an auxiliary receptor for BaEV (32) (33). Consequently, CD34+ cells are efficiently transduced with LV that are pseudotyped by either RD114 or BaEV envelope glycoproteins (34), particularly in the presence of VF1 (35) (36). Since syncytin 1 also uses ASCT1 and ASCT2 receptors for cell entry (15), it was expected that CD34+ cells would be transduced by LV-Syn1 in the presence of VF1. This was not the case, even though we used conditions previously tested in our laboratory (29) (26) and had 29±15% of cells expressing the transgene (n=9 donors) after one infection cycle with a LV-VSVg coding for GFP. LV-Syn1 in the presence of VF1 only led to 3±7% cell marking (n=8 donors). Practically no marking was obtained in the absence of VF1 (not shown). LV-Syn2 also failed to transduce CD34+ cells, achieving 0.1±0 marking (n=2 donors) in the presence or absence of VF1. The concentrations of vectors tested in these experiments were the same as in Figure 2, therefore contrasting the poor transduction of CD34+ cells with the efficient transduction of B and myeloid cells.

### Transduction of murine B cells in vitro and in vivo

Murine syncytin-pseudotyped LV, like their human counterparts, also transduced murine primary B cells *in vitro*. Without prior activation, spleen cells from C57Bl/6 mice were infected with either LV-SynA or LV-SynB in the presence of VF1 and high levels of transduction were obtained in CD19+B220+ cells after 4 days of culture, whereas CD3+ T cells were only transduced at moderate levels (Figure 3A) (gating strategy in supplementary Figure S3A). Transduction was confirmed by the detection of vector copies in the cells and the 2 murine syncytins had comparable effects (Figure 3B). Murine bone marrow (BM) cells were also transduced with LV-SynA or LV-SynB in the presence of VF1. After 4 days of culture, different stages of B cell maturation (pre-pro B cells, pro-B cells, pre-B cells and mature B cells) defined by differential expression of CD19, B220, CD43, CD24 (gating strategy shown in Supplementary Figure S3B) were transduced with similar efficiency by the 2 murine syncytins (Figure 3C). Transduction of murine BM cells was confirmed by integrated vector copies per cell (Figure 3D), although in these short-term experiments, some level of pseudo-transduction cannot be excluded.

**Fig. 3:**
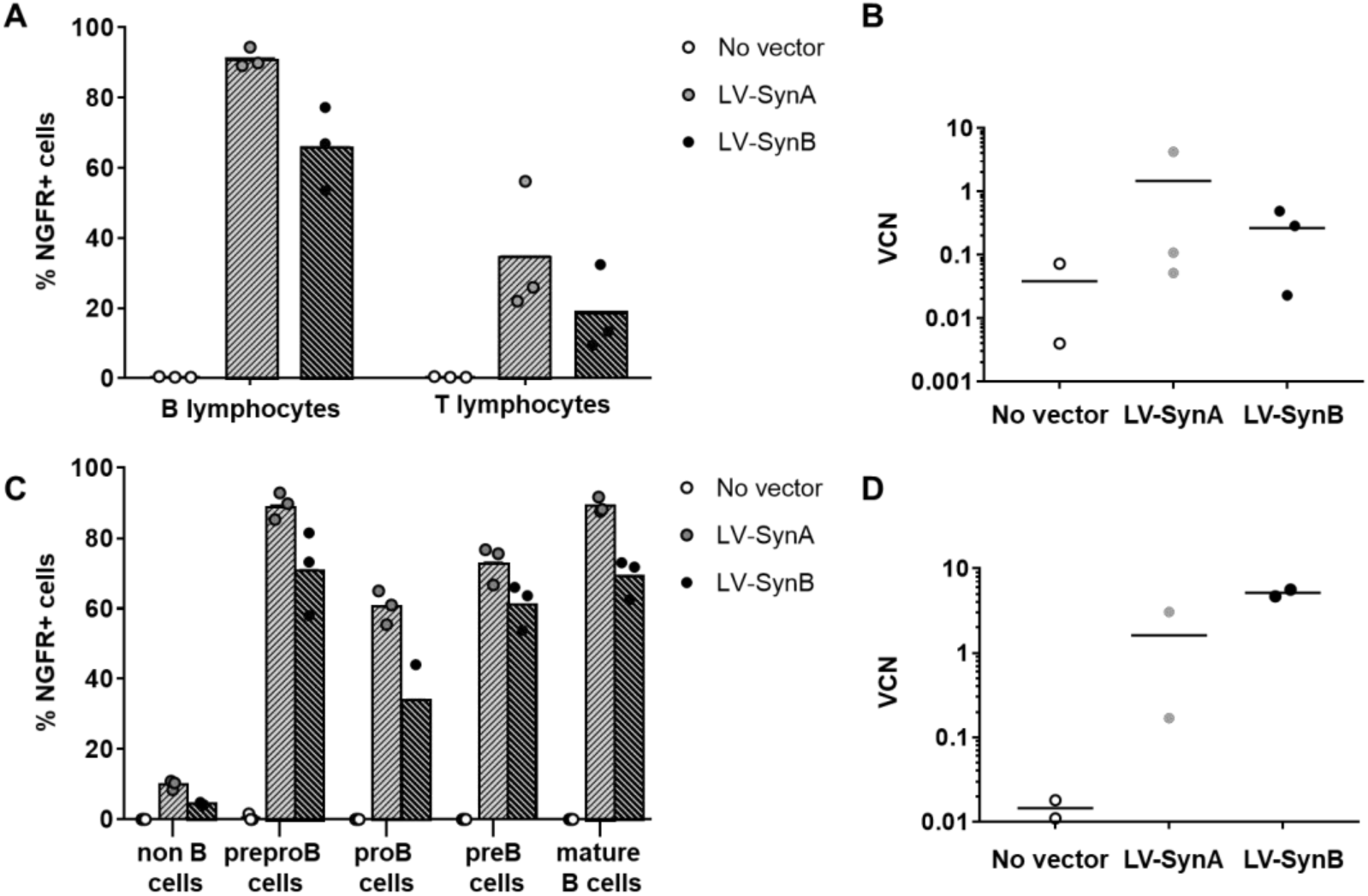
Transduction of primary B cells from spleen or from bone marrow with syncytin-pseudotyped vectors in mice. (**A-D**). Spleen cells (**A-B**) or bone marrow cells (**C-D**) from C57Bl/6 mice (n=3 mice) were infected with 1E+05 IG/mL LV-SynA or LV-SynB encoding NGFR in the presence of VF1 without prior activation and cells were cultured for 4 days and analyzed. (**A**) NGFR expression was measured by FACS in live spleen B cells (7AAD-CD19+B220+) and in live spleen T cells (7AAD-CD19-B220-CD3+). (**B**) Vector copy number in total spleen cells was determined by qPCR (B). (**C**) NGFR expression was measured by FACS in different subsets of bone marrow live cells (non B cells (7AAD-CD19-B220-), pre-pro-B cells (7AAD-CD19-B220+CD43+CD24-), pro-B cells (7AAD-CD19+B220+CD43+CD24-), pre-B cells (7AAD-CD19+B220+CD43lowCD24+) and mature B cells (7AAD-CD19+B220+CD43-CD24-)). (**D**) Vector copy number on total bone marrow cells was determined by qPCR.

The transduction of splenic B cells by LV-Syn vectors was confirmed *in vivo* in mice and over longer periods of time than *in vitro* studies. A single intravenous injection to mice of 5E+05 IG of LV-SynA expressing luciferase achieved stable gene expression in the spleen as visualized by a bioluminescence signal 3 weeks post-infusion (Figure 4A-B). The bioluminescent signal is significantly above background 7 days post infusion of LV-SynA and remains stable over time thereafter (Figure 4B). The transgene signal obtained after LV-SynA injection was not as high as that of LV-VSVg but it was notably more selective in terms of biodistribution, as it was not found in the liver contrary to VSVg (Figure 4A-B). The *in vivo* transduction of spleen by LV-SynA was confirmed by the detection of vector copies in whole spleen cells, 3 weeks post-infusion of the vector (Figure 4C). The levels were low but similar to those obtained following LV-VSVg injections. To ascertain that B cells were transduced, we sorted spleen CD19+ cells by FACS 3 weeks post-infusion of vector (purity > 93%). The presence of integrated vector could be observed by PCR in such purified CD19+ B cells prepared from different mice (Figure 4D), although the levels were too low to obtain reliable quantification. B cell transduction *in vivo* by LV-SynA was confirmed by immunohistological analysis of spleens indicating that follicular B220+ cells expressed the luciferase transgene (Figure 4E). It appears that cells other than B220+ cells are transduced *in vivo* as well, notably in the T cell area of the germinal centers, in coherence with *in vitro* results.

**Fig. 4:**
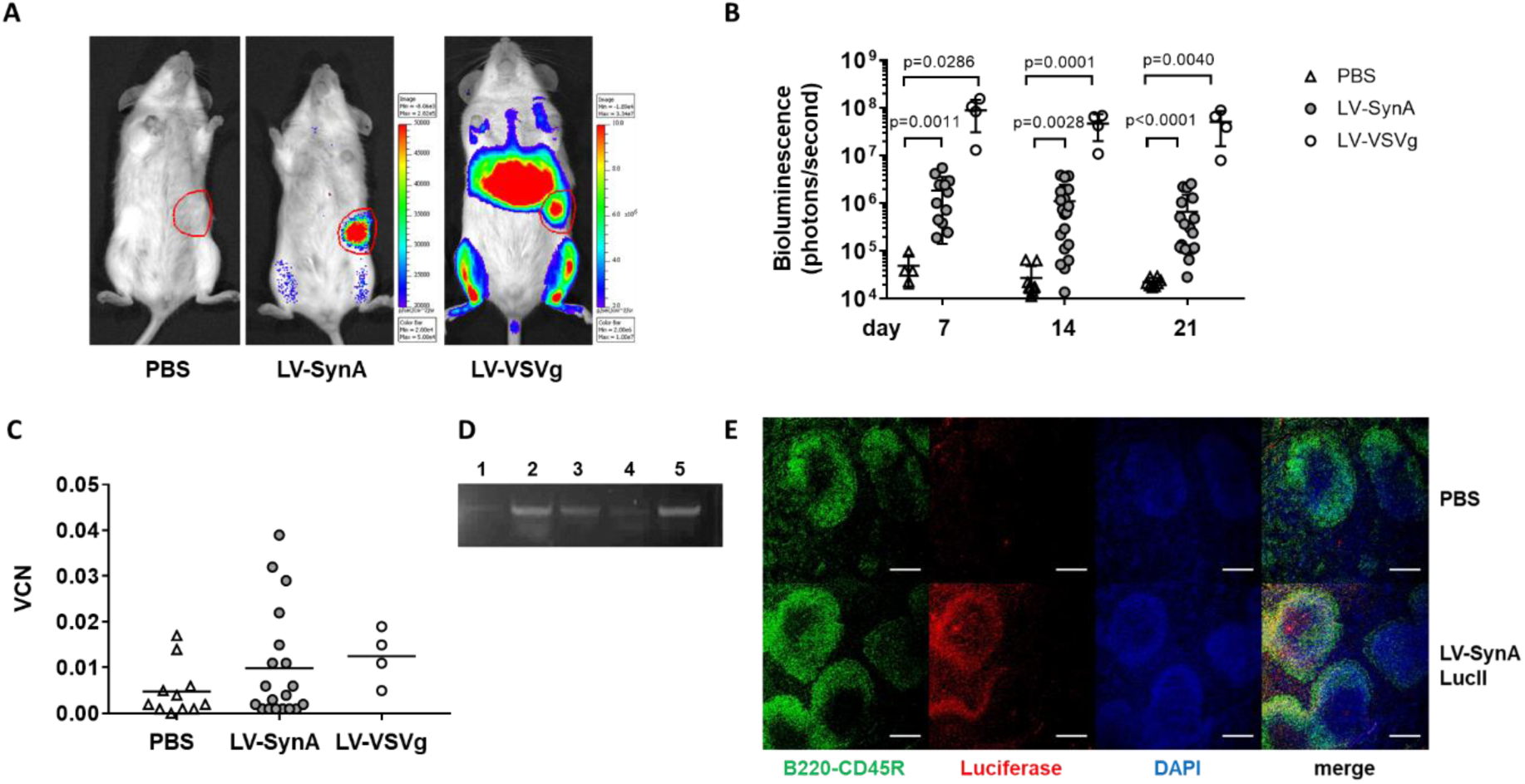
In vivo gene transfer into the spleen and into B cells following intravenous administration of LV-SynA vector. 6 week old C57Bl/6 albinos mice were injected intravenously with 5 E+05 IG per mouse of LV-SynA or LV-VSVg vectors encoding luciferase (LucII). The bioluminescent signal was measured over time and after 3 weeks, mice were sacrificed to confirm transduction and B cell gene transfer. (**A**) Representative results of bioluminescence in mice analyzed at 21 days. (11 mice tested in the PBS group, 18 mice injected with LV-SynA and 4 mice injected with LV-VSVg). The signals shown in the spleen of the PBS mouse is 2.3E+04 photons/second, in LV-SynA-injected mice 6.7E+05 photons/sec and in LV-VSVg-injected mice 5.1E+07 photons/second. (**B**) Bioluminescence was quantified in photon/second in the different groups over time. Each dot corresponds to one mouse analyzed. The signal in LV-SynA group was significant from the PBS. Statistics were calculated using a Mann-Whitney test. (**C**) Half of the spleen was used to determine vector copy number by qPCR and results for each mouse in the different groups are represented by a dot. (**D**) Half of the spleen was paraffin-embedded and sectionned to perform immunological staining with anti-B220 (CD45R) anti-luciferase and DAPI. Results of representative mice injected with PBS (top panel) or LV-SynA encoding LucII observed on the confocal microscope SP8 (Leica, Wetzlar, Germany) using the 10X objective (bottom panel). The bar represents 500 µm.

### Expression of the syncytin receptors on B cells

Possible interactions between B cells and syncytins have not yet been reported prompting us to search for the presence of syncytin receptors on human and murine B cells. ASCT2 and MFSD2a which are respectively the entry receptors for syncytin-1 and syncytin-2 (15, 17) were readily detected on CD19+ human blood B cells using indirect immunofluorescence and flow cytometry (Figure 5A). These receptors were also detected on CD11c+ dendritic/myeloid cells and at lower levels on CD3+ T cells as previously reported by others (37). Approximately 50% of blood CD19+ cells and of CD11c+ cells expressed detectable levels of ASCT2 and of MSFD2A whereas only about 10% of CD3+ T cells were positive (Figure 5B). The differential expression of these receptors in blood cell subsets was confirmed by qRT-PCR which confirmed mRNA expression in CD19+ B cells and the highest levels in CD11c+ cells (Figure 5C). Expression of ASCT2 and MFSD2a was also found on 293T cells whereas it was at a much lower level of HCT116 cells providing a likely explanation for their lack of transduction with the human LV-Syn vectors (Supplementary Figure S4). The expression of ASCT1 was not examined.

**Fig. 5:**
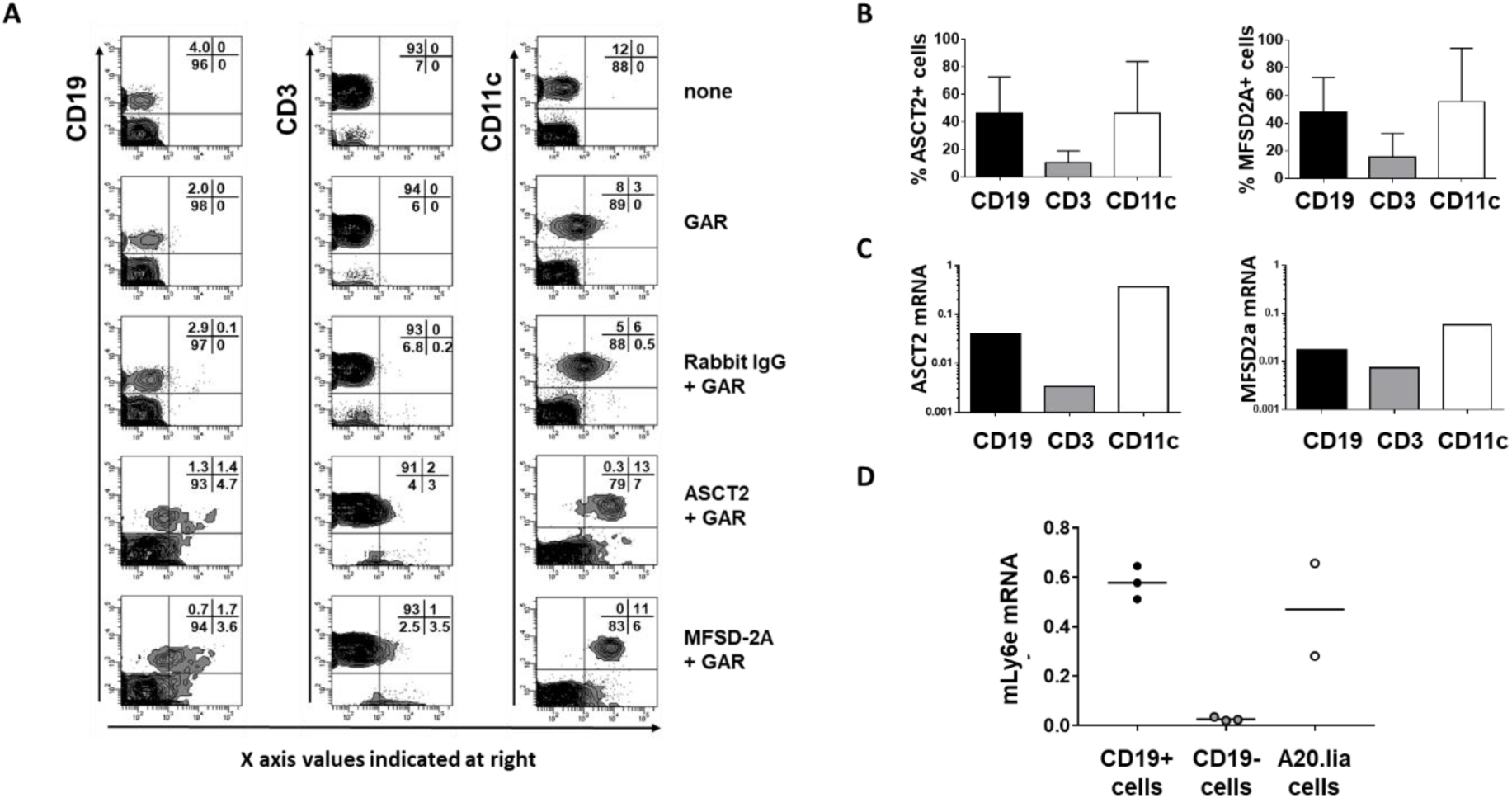
Expression of syncytin receptors on human and murine primary cells. (**A-C**) Analysis of ASCT2 and MFSD2a expression in human PBMCs. A. Representative FACS analysis. (**B**) Quantification of ASCT2 or MFSD2a protein expression by FACS (average and SD) in CD19+ B cells, CD3+ T cells and CD11c+ myeloid dendritic cells C. mRNA expression by qPCR in CD19+ B cells, CD3+ T cells and CD11c+ myeloid dendritic cells. Results from 3 experiments (1 blood donor each). (**C**) mRNA expression on mLy6e was measured by RT-qPCR on CD19+ or CD19-cells from mice spleens (3 different mice) and on A20.IIa cell line. Results are expressed in abundance of mLy6e mRNA.

The receptor for murine syncytin A was recently reported as Ly6e (19) and the receptor for the murine syncytin B has still not been identified. In the absence of antibodies suitable for flow cytometry, we used qRT-PCR to demonstrate higher expression of Ly6e mRNA in murine spleen sorted CD19+ B cells, whereas the CD19-cell population expressed low levels of mRNA (Figure 5D). Equivalent levels of Ly6e mRNA were found in primary CD19+B cells and in the A20.IIa cell line. These results are coherent with the ability to transduce these cells with syncytin A-pseudotyped LV.

### Lower immune responses induced by syncytin-pseudotyped vectors in vivo

The administration of viral gene transfer vectors *in vivo* can induce complex adaptive immune responses dominated by immune recognition of the viral components of vectors and in some cases, by a response against the transgene product. The intravenous administration of LV-VSVg to mice is known to immunize against the transgene (5) linked in part to the interaction of the LV-VSVg particles with professional phagocytes in the liver (7). The intrinsic immunogenicity of LV is less known. We found that a single intravenous injection of LV-VSVg to mice induced robust antibody responses against the p24 capsid in mice (Figure 6). Replacing the viral glycoproteins of the envelope by syncytin A which is an endogenous cellular protein in mice, induced much lower levels of antibodies against p24 compared to LV-VSVg for the same amount of particles injected.

**Fig. 6:**
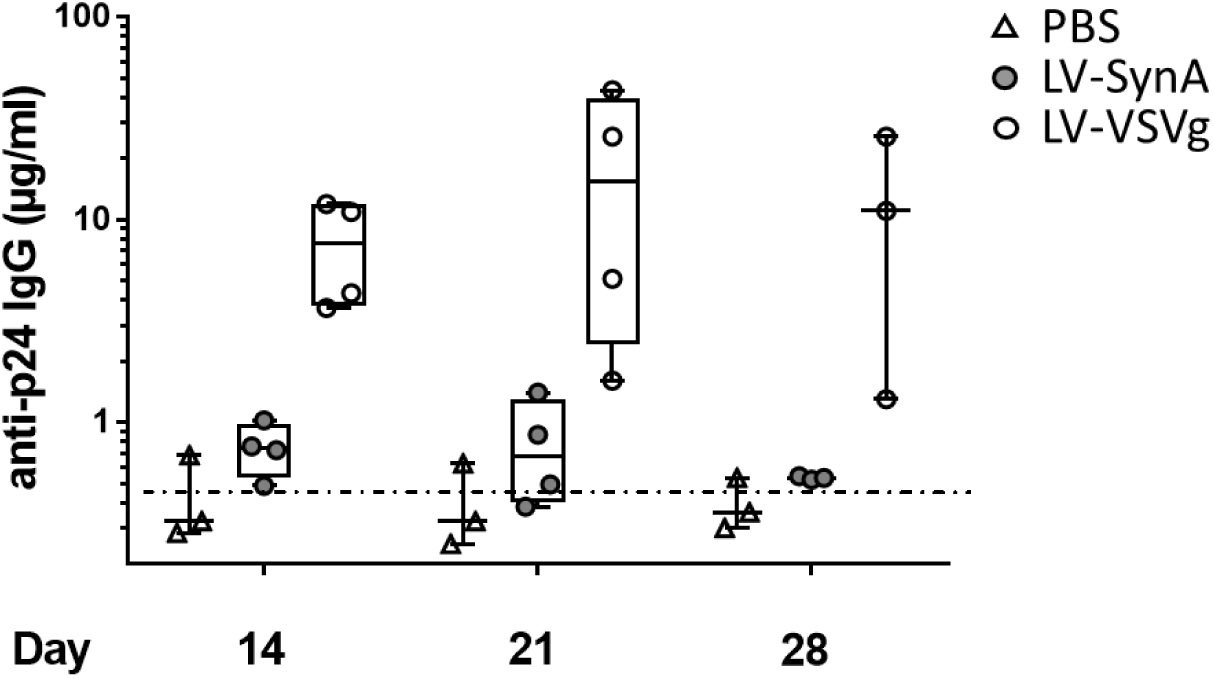
Low induction of anti-P24 antibodies by Lv-SynA administration contrary to LV-VSVg. C57BL/6 mice (n=4 in each group) were injected in the tail vein with either PBS or with the same amount of physical particles of LV-SynA or of LV-VSVg each encoding the same GFP cassette (injections of 4E+09 pp of vector /mouse representing similar values of p24 which were 4200ng p24 for LV-VSVg and 3700ng p24 for LV-SynA). Blood was sampled at different time points following vector administration and the presence of antibodies to p24 was measured by ELISA and expressed as µg/mL against a mIgG standard. The ELISA detection threshold was 0.25 µg/mL.

## Discussion

This paper shows for the first time that human and murine primary B cells functionally interact with syncytins because B cells are transduced efficiently by syncytin-pseudotyped LVs and B cells express known syncytin cellular receptors. Human B cells express high levels of ASCT2 and MFSD2a, receptors respectively for syncytin-1 and syncytin-2 (16, 17). Murine B cells express mLy6e mRNA, the recently-identified receptor for murine syncytin A (19). Previous reports in the litteratire showed that cultured human myeloid dendritic cells, and to a lesser extent T cells, expressed ASCT2 and MFSD2a (37) but B cells had not been examined in that study. We observed similar levels of ASCT2 and MFSD2a receptors in B cells and CD11c+ dendritic cells and accordingly, both of these cell subsets could be transduced at high levels with either of the LV-Syn1 or LV-Syn2 vectors. At this stage, we describe a functional interaction between syncytins and B cells but cannot exclude that other receptors or accessory molecules are involved in B cells:syncytin interactions. Nevertheless this offers novel possibilities for gene transfer into B cells.

These findings shed a novel light on the interactions of syncytins with the immune system and on the interactions between endogenous retroviral glycoproteins and B cells. Syncytins are involved in somatic cell fusion notably the formation of the syncytiotrophoblast layer in the placenta (13), myoblast fusion during skeletal muscle regeneration (20, 21) and the formation of multinucleated osteoclasts (22). While it is not clear how B cells relate to somatic cell:cell fusion processes, B cells may play a role in another function of syncytins which is their immunoregulatory effects. Retroviral envelopes carry an immunosuppressive peptide sequence in their transmembrane chain and this motif is also found in syncytins (23). The mechanism of action of this peptide sequence remains unclear. It has been postulated to involve antigen-presenting cells by modulating dendritic cell-mediated T cell activation (37) or could directed reduce the activation of T cells as shown by peptide-treated T cell lines (38). Our results suggest that B cells could potentially be involved and further studies are warranted on this aspect. Physiologically, B cells have the possibility to interact with syncytin-expressing cells or with syncytin-bearing particles at least during pregancy. Syncytins are found in the syncytiotrophoblast layer of placenta that mediates immune tolerance during pregnancy by separating the fetal and maternal blood circulation. Additionally, placental exosomes bearing syncytins are known to circulate and could mediate long-range interactions between syncytins and B cells. Whether or not syncytins modulate B cell function is not clearly obvious as proliferation and differentiation of human B cells was not affected, but more subtle mechanisms could be involved and deserve further investigation. Apart from a putative immunomodulatory effect on immune cells, syncytins could simply but very effectively render the LV particles invisible from immune recognition by endogenizing the envelope. Our results confirm that *in vivo* administration of LV-VSVG can be immunogenic and show for the first time that a robust humoral immune responses is induced against p24 a capsid component of LV following systemic administration to immunocompetent mice. Antibodies to the capsid could prevent further administration of the vector or may exacerbate immune rejection of transduced cells and could contribute to the loss of transgene expression following the use of LV-VSVg, as observed by others (7). We also confirm the strong tropism of LV-VSVG to liver already reported (5, 7) and show that in striking contrast, LV-SynA particles do not target the liver. Thus, as postulated by others, the envelope glycoproteins strongly determine tissue tropism and immune recognition of the particles and our results confirm that such glycoproteins can be engineered to change these interactions in a significant manner. Thus, further study is warranted to understand the precise mechanisms involved in syncytin-mediated tolerance to vector administration and B cell interactions. Interestingly, all syncytins studied as LV pseudotype have demonstrated an ability to transduce B cells and this is also the case for another endogenous envelope, the BaEV glycoprotein which allows high levels of gene transfer into activated or resting human B cells, at similar levels as observed with syncytins (39). The BaEV glycoprotein shares the same cellular entry receptors ASCT1 and ASCT2 as the human syncytin-1 (15) therefore our results showing ASCT2 exression on human B cells provides an explanation for the B cell tropism of BaEV. Other similarities between BaEV and syncytins include the need for a transduction additive when using these glycoproteins as LV pseudotypes (36). However, contrary to BaEV, syncytins do not mediate efficient T cell transduction or CD34+ cell transduction (40). Thus, different levels of receptors or other mechanisms such as entry or post-entry restriction steps may explain the differences between BaEV and syncytin cellular tropism.

Whereas B cells are essential players in the immune system, specific tools are required for efficient gene transfer in primary B cells either for cellular studies or to develop therapeutic approaches. The commonly-used VSVg-pseudotyped vectors are inefficient due to the lack of the LDL receptor expression on resting B cells (4). Besides natural pseudotypes such as BaEV (4), B cell-targeting LV systems have been engineered and were reviewed recently (8). Human B cells can be transduced by LV displaying Edmonston strain measles virus (MV) glycoproteins using either the native H/F capacity (41) or engineering them with mutations and a single chain Fv (ScFv) targeting hCD20 (42) or hCD19 (43). These MV-based systems enable about 20-40% transduction of activated peripheral blood human B cells, including malignant activated cells. Thus native syncytin glycoproteins are at least as efficient as MV-LV being capable of transducing almost all peripheral blood B cells with high concentrations of syncytin-LV. However, syncytins are distinct from MV as they did not efficiently mediate the transduction of established EBV-transformed B-LCL cell lines or of B cells during EBV transformation even though a few stably transduced cells could be obtained (not shown). Another unique feature of syncytins is their cross-species ability, since human syncytins can enable gene transfer into murine cells and murine syncytins enable gene transfer into human B cells, which is not the case for other non-VSVG envelopes such as GaLV, BaEV, RD114 or engineered MV-LV. Cross-species ability is advantageous in experimental systems especially for preclinical studies in mice.

Our results show an efficient transduction of B cells *in vitro* and *in vivo* but we do not claim that syncytins provide B-cell-restricted tropism. Several tissues are known to express receptors for syncytins but have not been studied yet. Placental layers express very high levels of MFSD2a (17). MFSD2a is also found and specifically localized in the microvessels of the murine blood brain barrier where it plays an essential role during brain development (44) but it is not known if these tissues would be targeted by vectors for gene delivery. Thus, the display of various human or murine syncytins on gene transfer vectors provides an interesting new tool for *in vivo* studies and possibly to envision new therapeutic developments but in-depth bioditribution studies should be conducted to determine the specificity of gene deliverey with these tools. Further optimization of vector titers will also be needed for efficient gene delivery in vivo. Certainly these tools could offer an interesting concept to avoid immune recognition which is currently a main limitation of gene therapy.

The transduction additive VF1 revealed the infectivity of syncytin vectors which was inapparent *in vitro* without this agent, but VF1 was not required *in vivo* and this difference sheds some light on the mode of action of VF1. This peptide is a short histidine-rich, amphipathic peptide derived from the LAH4 family of DNA transfection agents (26, 28) (27). This family of peptides is able to bind to nucleic acids and to disrupt endosomal membranes when activated at low pH enabling them to deliver their cargo efficiently and with low associated toxicity (45). Within this family, VF1 is a specific derivative with lysine residues at the N-terminal extremity containing leucine hydrophobic residues and with a specific hydrophilic angle formed by histidine residues that are important for its activity. VF1 is an efficient retroviral and lentiviral transduction enhancer with pseudotypes such as GALV, GALV-TR, RD114-TR or MLV-A as well as with VSVg which all use receptor-mediated endosomal entry. VF1 enhances LV transduction of target cells by enhancing the adhesion and fusion steps (46) and by promoting viral pull-down following formation of nanofibrils in medium (47). It is therefore interesting that VF1 also promotes the transduction of target cells mediated by syncytins considering the high fusogenic activity of these proteins, which we observed during production and which is not observed to this level with other glycoprotein LV pseudotypes. Thus, it is likely that VF1 manily enhances transduction through the facilitation of vector-cell contact. This would explain why the addition of VF1 is not necessary *in vivo*, as contacts between syncytin-LV and target cells may occur in more restricted spatial environment as in culture dishes.

Altogether, our findings contribute to a better understanding of host:vector interactions with possible impact in immunology and gene therapy. The concept of endogenizing the surface of the gene transfer particles with cellular fusion proteins instead of viral glycoproteins could facilitate the use of LV gene transfer *in vivo*. Our findings also opens possible new applications to use synyctins to target B cells in immunotherapy of cancer, transplantation, genetic or infectious diseases and in biotechnology engineering.

## Materials and Methods

### Ethical and regulatory aspects

Studies in mice and with human cells were conducted following international ethical guidelines and were authorized by the french ministry of higher education research and innovation with: approval for studies in mice APAFIS#3744-2016012210018523v2 - ethical committee 51; agreement n° 3664 for the use of GMO and declaration DC-2018-3276 for the use of human samples for research.

### Animals

C57BL/6 mice and C57BL/6 albinos (B6N-Tyrc-Brd/BrdCrCrl) 6 week old mice (Charles River, Italy) were housed in SPF conditions at the CERFE facility (Evry) operated by Charles River. For injections, 100µL volume was used for intravenous (IV) (tail vein) or intraperitoneal (IP) routes and mice were anesthetized with ketamine (120mg/kg) and xylazine (6mg/kg).

### Human and murine cells

Human blood of healthy volunteers was purchased from the french blood bank (EFS, Evry, France). Umbilical cord blood was obtained with consent from uncomplicated birth at the Centre Hospitalier Sud-Francilien (CHSF) in Evry. Human embryonic kidney 293T cells (Genethon), human colorectal carcinoma HCT116 cells (CCL-247; ATCC, Manassas, VA), murine embryonic fibroblast NIH/3T3 cells (CRL-1658, ATCC) and murine myoblasts C2C12 (CRL-1772, ATCC) were cultured at 37°C, 5% CO2 in Dulbecco’s modified Eagle’s medium (DMEM) supplemented with Glutamax and 10% of heat-inactivated fetal calf serum (FCS) (Life Technologies, St-Aubin, France). Human choriocarcinoma BeWO cells (CCL-98; ATCC) were cultured in Ham’s F12K medium (Life technologies) supplemented with 10% of FCS. Raji cells, Jurkat cells were cultured in XVivo 20 medium (Lonza, Levallois-Perret, France). A20.IIa murine lymphoma cells (cellosaurus A20.IIa (RRID:CVCL_0J27), kindly provided by S. Fisson) were cultured in Roswell Park Memorial Institute medium (RPMI) supplemented with 10% of heat inactivated FCS, 1% of Glutamax, 1mM of sodium pyruvate (Life Technologies) and 50µM of β-mercaptoethanol (Sigma Aldrich).

### Cloning of syncytin-expressing plasmids

pcDNA3.1-syncytin-1, pcDNA3.1-syncytin-2, pcDNA3.1-syncytin-A, pcDNA3.1-syncytin-B plasmids were constructed by cloning synthesized DNA sequences (Gencust, Dudelange, Luxembourg) (respectively Ensembl genome browser ENST00000493463, ENSG00000244476, ENSMUSG00000085957 and ENSMUSG00000047977 transcript sequences), into a pcDNA3.1 plasmid (Invitrogen, Carlsbad, California) using standard cloning techniques. Plasmids were produced with endotoxin-free reagents using DNA/RNA purification Nucleobond kit (Macherey-Nagel, Duren, Germany). Plasmid sequences were verified by two strand sequencing.

### Production and titration of vectors

Lentiviral particles pseudotyped with either syncytin-1, syncytin-2, syncytin-A or syncytin-B were produced by transient transfection of 293T cells with 4 plasmids, using calcium phosphate. Each T175 cm2 flask was transfected with 14.6 µg pKLgagpol expressing the HIV-1 gagpol gene, 5.6 µg pKRev expressing HIV-1 rev, 20µg of either one of the syncytin envelope plasmids and 22.5 µg of the gene transfer lentiviral cassette (pCCLsincPPT-PGKeGFP-WPRE expressing enhanced green fluorescent protein (GFP) under the control of the human phospho glycerate kinase (PGK) promoter; or pRRLsincPPT-PGKdelta-NGFR-WPRE expressing a truncated form of the nerve growth factor receptor (NGFR) under the control of the human PGK promoter; or pRRLsincPPT-SFFV-LucII-WPRE expressing luciferase II under the control of the spleen focus-forming virus (SFFV) long-terminal repeat (LTR) promoter). Medium was changed the day following transfection and replaced by DMEM 4.5g/L glucose supplemented with penicillin and streptomycin and 10% FCS. After 24 hours, medium containing viral particles was collected, clarified by low speed centrifugation and filtration through 0.45µm filter, then concentrated by ultracentrifugation at 50 000g for 2h at 12°C. The pellet was resuspended in PBS, aliquoted and stored at −80°C.

VSVg-pseudotype LV were produced by transient transfection as previously described (26). Physical particle titers of VSVg- or syncytin-pseudotyped LV were determined by p24 ELISA (Alliance© HIV-1 Elisa kit, Perkin-Elmer, Villebon/Yvette, france) and by Nanoparticle Tracking Analysis with the Nanosight NS300 (Malvern Panalytical, Orsay, France). Infectious titers were determined by testing serial dilutions of vectors on A20.IIa cells in the presence of the peptide Vectofusin-1 (VF1) prepared as described (26) and used at a final concentration of 12µg/mL. After 7 days of culture, A20.IIa cells were analyzed by qPCR to measure infectious genomes (IG)/mL using standard calculations (48). In some experiments and as indicated, infectious titers of GFP-encoding vectors were also determined as transducing units (TU)/mL in 293T cells using FACS.

### Culture and transduction of human peripheral blood mononuclear cells

Human peripheral blood mononuclear cells (PBMC) obtained by Ficoll gradient centrifugation (Eurobio, les Ulis, France) were suspended in X-VIVO 20 serum-free medium for transduction. When indicated, IL-7 (10ng/mL) (Miltenyi Biotech, Bergisch Gladbach, Germany) was added prior to transduction. Cells were infected for 6 hours with LV in the presence of VF1 (12µg/mL), then washed and X-VIVO 20 supplemented with 10% FCS and IL-7 (10 ng/mL) was added and cells were cultured for up to 7 days. To expand B cells, CD40L (2 µg/mL) and IL-21 (50 ng/mL) were added to the culture medium and cells were cultured up to 2 weeks. For proliferation studies, PBMC were labelled with 2µM CFSE (Carboxy Fluorescein Succinimidyl Ester) (Molecular Probes, Cambridge, UK).

### Culture and transduction of murine primary cells

Red blood cell-depleted leukocytes were prepared by shredding the spleen of C57BL/6 mice on a 70µm filter in RPMI medium or from the femurs and tibias flushed with RPMI medium, followed by red cell lysis using ACK buffer. The transduction of either spleen or bone marrow cells was done by culturing 2E+05 cells in 96 wells plates in RPMI medium supplemented with 10% FCS, 1% Glutamax and 50µM of β-mercaptoethanol and adding immediately in the wells LV-SynA or LV-SynB (1E+05 IG/mL) and 12µg/mL of VF1 in a final volume of 200µL. After 6 hours, cells were collected, centrifuged (500g), the medium was changed and cells were cultured for 4 days in the same medium.

### Flow cytometry (FACS)

Antibodies used are listed in supplementary table S2. Prior to adding antibodies to cells, Fc receptors were blocked with either gammaglobulin (1mg/mL) (Sigma Aldrich) for human cells or with CD16/CD32 antibodies for murine cells, during 5 min. at 4°C. Staining was done with saturating amounts of antibodies for 30 min. at 4°C in PBS with 0.1% bovine serum albumin (BSA) (Sigma Aldrich), cells were washed twice and 7-amino-actinomycin D (0.3 mg/mL) (Sigma-Aldrich) was added to enable dead cell exclusion. The LSRII cytometer using Diva software (BD-Biosciences) was used and data were analysed with Kaluza (Beckman-Coulter, Villepinte, France) or FlowJo v10 (BD-Biosciences) softwares.

### In vivo luciferase imaging

Bioluminescence imaging of mice was started 10 min after IP injection of D-luciferin (150µg/mL) (Interchim) using a CCD camera ISO14N4191 (IVIS Lumina, Xenogen, MA, USA). A 3 min bioluminescent image was obtained using 10 cm field-of-view, binning (resolution) factor 4, 1/f stop and open filter. Regions of interest (ROIs) were defined manually (using a standard area), signal intensities were calculated using the living image 3.2 software (Xenogen) and expressed as photons per second. Background photon flux was defined from an ROI drawn over the control mice in which no vector had been administered.

### Murine B cell sorting and PCR

Immediately after sacrifice of mice, one half of each spleen was perfused with collagenase IV (1mg/mL, Invitrogen) and DNase I (50µg/mL, Roche) and then incubated at 37°C 45 min to dislodge as many cells as possible. The reaction was stopped by the addition of EDTA (100mM), cells were isolated by mechanical shredding and stained with anti-B220, anti-CD19, anti-CD3 and anti-CD11b antibodies (antibodies listed in supplementary Table S2), washed in PBS with 0.1% BSA and sorted by FACS on the MoFlo® Astrios (Beckman Coulter).

Genomic DNA was extracted from the sorted subpopulations of B cells (CD19+B220+CD3-CD11b-) and T cells (CD19-B220-CD3+CD11b-) using Nucleospin Tissue kit (Macherey-Nagel). Provirus was amplified by 35 cycles of PCR using the PSI_F 5’ and WPRE_R primers. PCR primer sequences are listed in supplementary Table S3.

### Immunohistofluorescence on murine spleen

After sacrifice, one half of the spleen of each mouse was paraffine-embedded for immunohistology. Microtome sections (4µm) were saturated with PBS-BSA 5% during 30 min. at room temperature. Sections were stained with a polyclonal antibody anti-luciferase (Novus biologicals) diluted at 1/100 and an antibody anti-CD45R/B220 (BD Pharmingen) diluted at 1/20 as primary antibodies, a donkey anti-goat AlexaFluor 594 (Invitrogen) diluted at 1/1000 and a goat anti-rat AlexaFluor 488 (Invitrogen) diluted at 1/600 as secondary antibodies. Primary antibodies were incubated overnight at 4°C in a humidity chamber and secondary antibodies were incubated for 90 min at room temperature.

### Receptor studies

In human cells, total RNA was extracted (SV total RNA isolation, Promega, Charbonnière les bains, France) from CD19+ CD3-CD11c-cells, CD3+ CD19+CD11c-cells and CD11c+ CD3-CD19-cells sorted by FACS (> 95% purity) from PBMC. Residual DNA was removed using the free DNA kit (Ambion, Courtaboeuf, France), cDNA was synthesized using random hexamers (Superscript II first strand synthesis system, Invitrogen). Real time PCR (7900HT, AB Biosystems) was used to quantify either ASCT2 or MFSD2A relative to the housekeeping gene TFIID using primers and probes listed in supplementary Table S3. The relative abundance was calculated using the formula: relative abundance = 2^-ΔΔCt with ΔCt= Ct gene of interest – Ct housekeeping gene and ΔΔCt= ΔCtsample – ΔCtcalibrator.

RNA was extracted from murine CD19+ and CD19-spleen B cells which were separated by magnetic bead selection (Miltenyi) using the RNeasy mini kit (Qiagen). Reverse transcription was performed with random hexamers using Verso cDNA Synthesis Kit (Invitrogen). Real time PCR was performed using LightCycler 480 system (Roche, Basel, Switzerland) to amplify mLy6e with primers listed in Supplementary Table S3 and Sybr Green PCR Master Mix (Applied Biosystem, Foster city, CA, USA). Results are expressed in relative abundance to the P0 housekeeping gene.

### Detection of murine anti-p24 antibodies by ELISA

Recombinant p24 (Abcam, Cambridge, U.K.) was bound to plates using high binding 96-well Nunc MaxiSorp Immunoplate (Thermo Fisher Scientific) coated 2 hours at 37 °C with 100 ng per well of p24 recombinant protein diluted at 1 µg/mL in 0.2 M carbonate/bicarbonate buffer in pH 9.5, then wells were blocked with PBS-BSA 2% for 2 h at room temperature. A mouse anti-HIV p24-specific monoclonal antibody (clone 38/8.7.47) (Abcam, Cambridge, U.K.) was used as standard and tested in parallel with serum samples diluted in assay buffer (PBS BSA 0.5%). Samples were incubated 2 hours at room temperature. After 3 washes, HRP-conjugated polyclonal goat anti-mouse IgG antibodies (Jackson Immuno Research, Newmarket, U.K.) (1/1000 dilution) were added, incubated 1h at room temperature and reactions were revealed by adding 3,3′,5,5′-tetramethylbenzidine substrate (TMB) (BD Biosciences) and measuring OD at 450 and 570 nm (for background substraction) (Enspire plate reader, Perkin Elmer) after blocking the reaction with H2SO4.

### Statistical analysis

Statistical significance was assessed as specified, using GraphPad Prism software (GraphPad Inc, La Jolla, CA).

## Supplementary Materials

### Supplementary figures

**Fig. S1:**
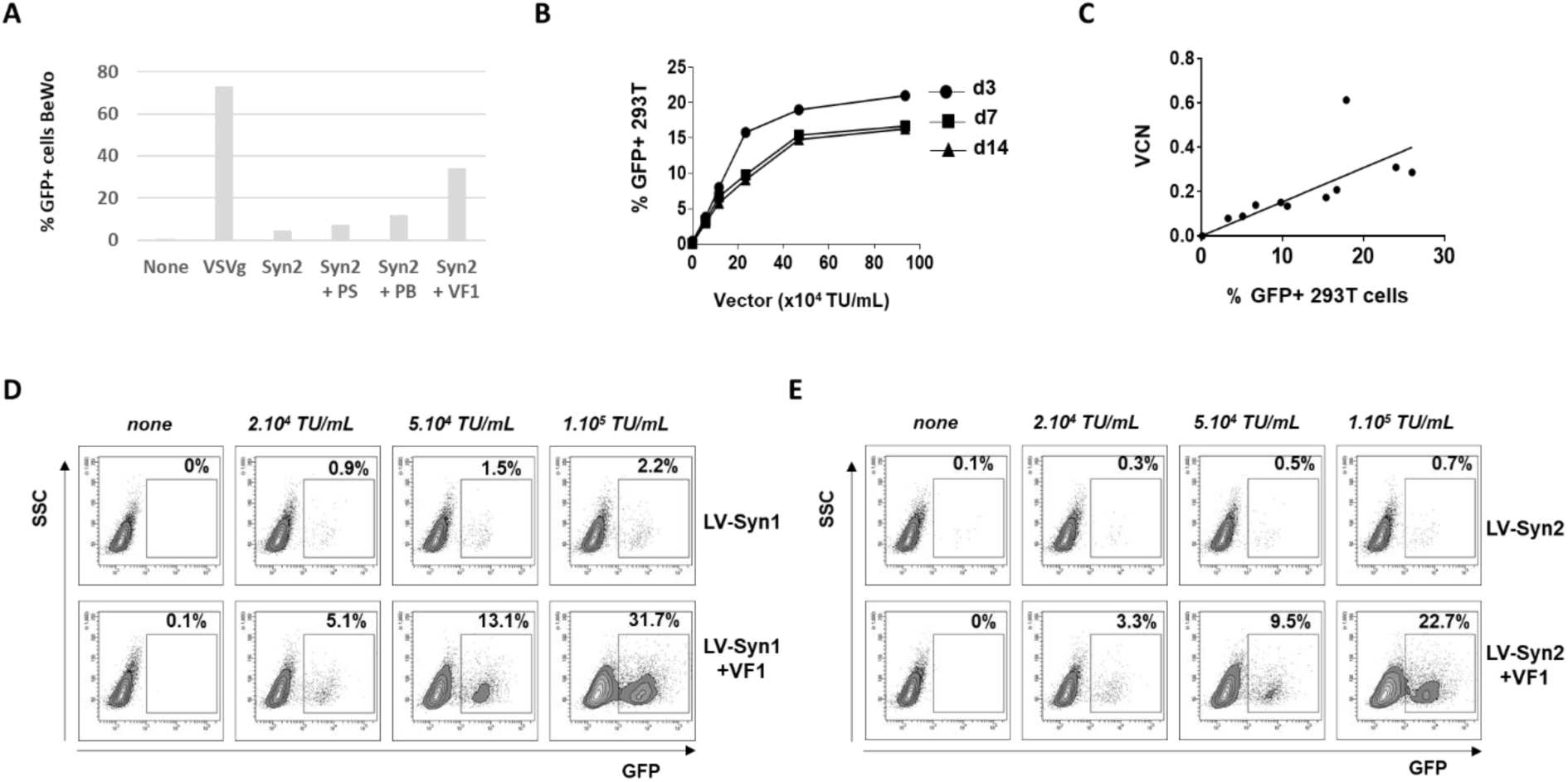
Gene transfer with LV-Syn vectors is enhanced by VF1, stable over time and vector dose-dependent. A. Transduction of BeWO cells with LV-Syn2 vector encoding GFP (1E+05 TU/mL). In these experiments, protamine sulfate (PS) (Sigma-Aldrich, St Louis, MO) and polybrene (PB) (Sigma-Aldrich) were respectively used at final concentrations of 8µg/mL and 6 µg/mL in the transduction culture medium. B. Dose-dependent increase in transduction of 293T cells and stability of transduction over time. 293T cells were transduced with increasing concentrations of LV-Syn1 encoding GFP (concentrations based on transducing units (TU)/mL defined by FACS titration on 293T cells in the presence of VF1). Cells were cultured for the indicated times (d=days) at which GFP expression was measured by FACS. Results are expressed as percentage of GFP+ cells in the culture. Results of 1 experiment representative of 4. C. Cells transduced as in (B) were tested at day 3 to correlate GFP expression and proviral integration with quantification of vector copy number per cell (VCN) by qPCR. D-E. Representative FACS plot showing the dose-dependent transduction of 293T cells. Increasing concentrations of LV-Syn1 (E) or LV-Syn2 (F) expressing GFP (titered as in B) were used to transduce 293T cells in the absence or in the presence of VF1. FACS analysis was performed at day 3. Results are representative of 5 separate experiments.

**Fig. S2:**
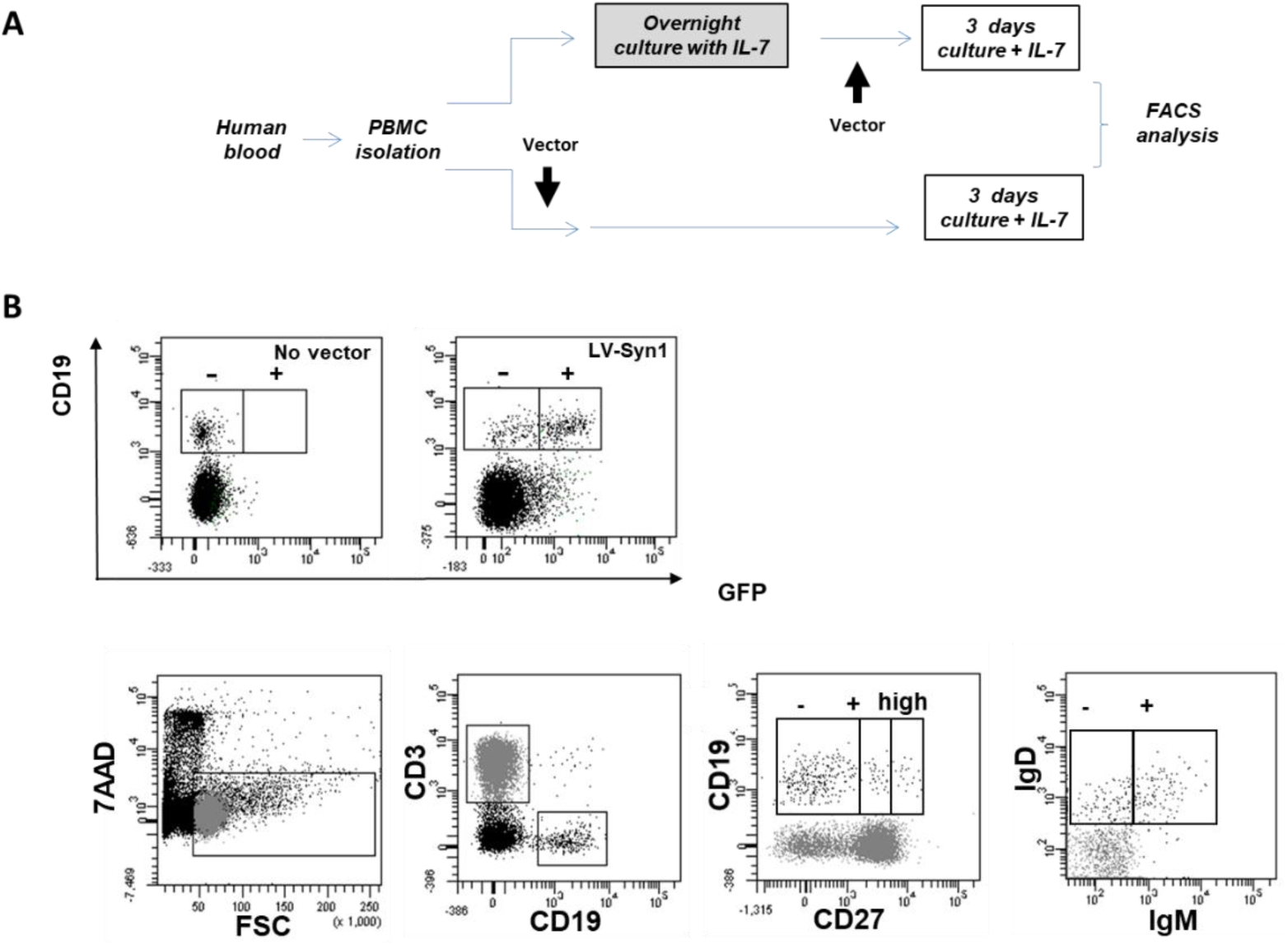
Experimental schema and gating strategy for human PBMC analyses. A. Treatment schema for infection of PBMC with LV-Syn. B. FACS gating strategy to analyse the transduction of various B cell subpopulations

**Fig. S3:**
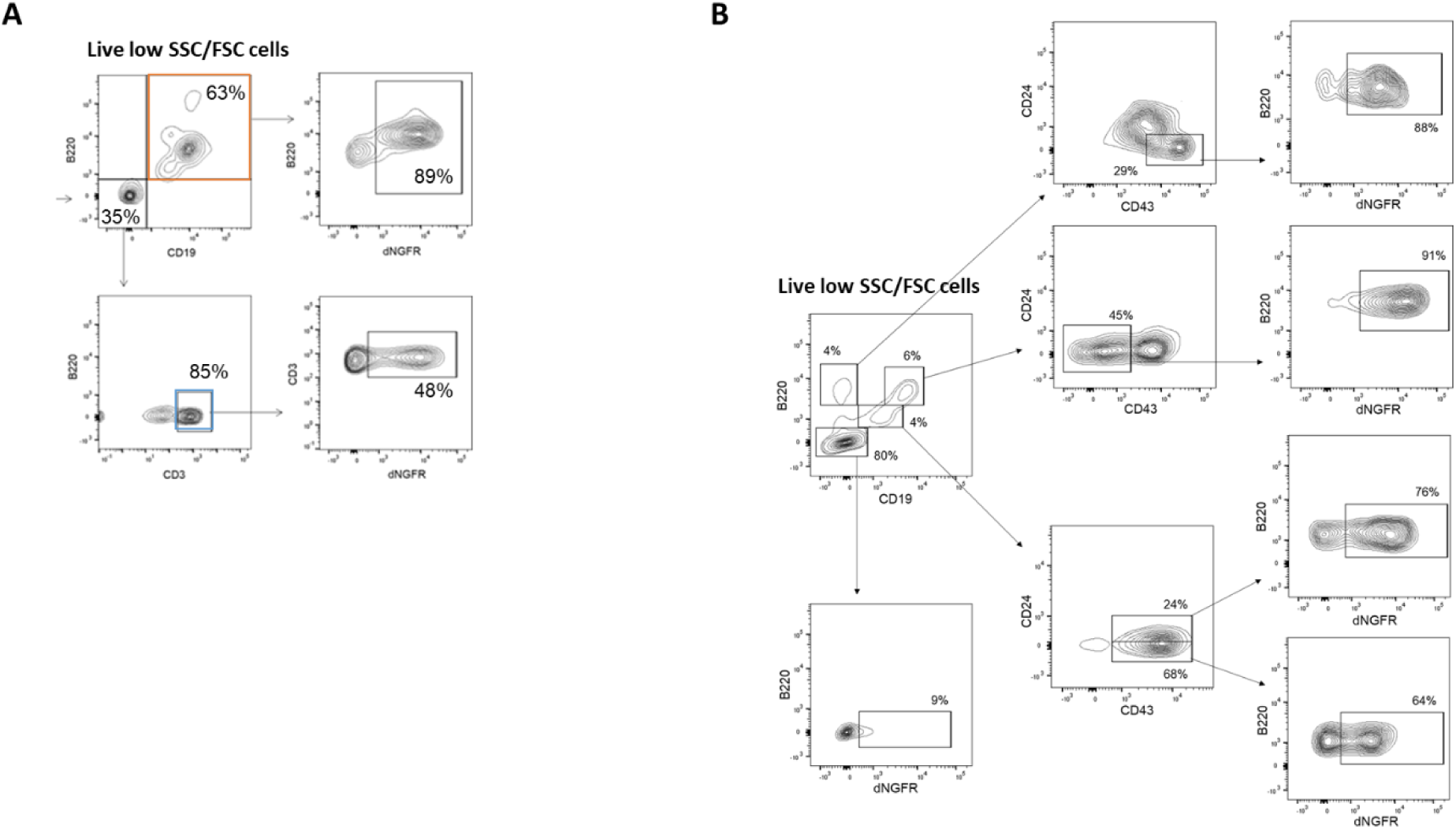
Experimental schema and gating strategy for murine analyses. A. FACS gating strategy to analyze the transduction of spleen B and T cells B. FACS gating strategy to analyse the transduction of bone marrow cells.

**Fig. S4:**
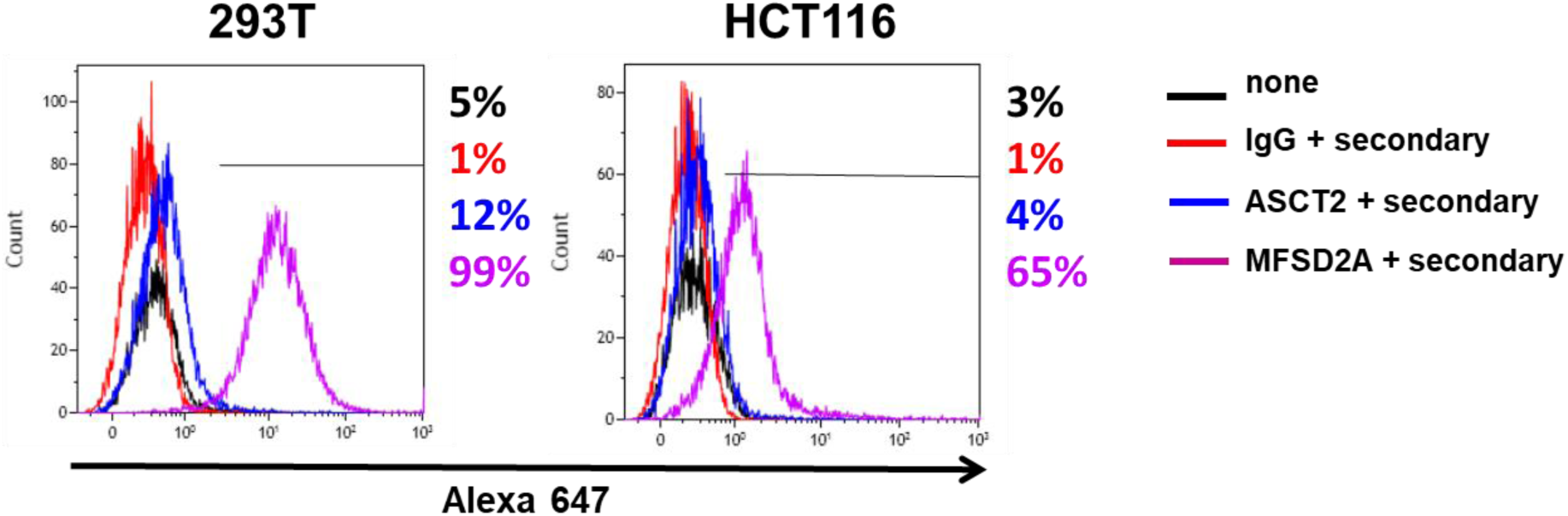
Receptor expression on human cell lines. Immunostaining for ASCT2 and MFSD2a on HEK293T cells and HCT116 cells. Histograms represent the mean fluorescence intensity (MFI) of unstained negative control cells (black), cells stained with irrelevant IgG and the Alexa 647-conjugated secondary antibody (red), cells stained with the anti-ASCT2 and secondary antibody (blue) and cells stained with anti-MFSD2a and secondary antibody (purple).

### Supplementary tables

**Table S1:** Transduction of PBMC with LV-Syn1 and LV-Syn2 in the presence of VF-1. Legend : Average percentage ± SD of total nucleated cells (CD45+) expressing GFP, 3 days after infection of PBMC. The cells were preactivated with IL-7 or not as indicated, and were transduced with indicated LV in the presence or absence of Vectofusin-1 (VF1). PBMC were obtained from different donors and the number of donors (n) tested is indicated. Data obtained from 3 experiments without VF1 and 7 to 9 experiments with VF1. (*) p<0.05 compared to “no vector” in a paired Student’s t test

|  | No vector |  | LV-Syn1 |  | LV-Syn2 |  | LV-VSVg |  |
| --- | --- | --- | --- | --- | --- | --- | --- | --- |
| VF1 | - | + | - | + | - | + | - | + |
| Prior activation with IL-7 |  |  |  |  |  |  |  |  |
| **Average ± SD** | 0.1±0.1 | 0.3±0.2 | 1.0±0.1 | **12.7±14.6\*** | 1.7±1 | **4.5±2.7\*** | **5.9±7.8\*** | **5.4±1\*** |
| **n** | 9 | 7 | 3 | 9 | 3 | 7 | 9 | 6 |
| Non-activated PBMC |  |  |  |  |  |  |  |  |
| **Average ± SD** | 0.2±0.2 | 0.7±0.7 | 2.7±3.3 | **5.7±6.7\*** | 1.4±1.6 | 0.8±0.5 | **4.0±2.0\*** | **6.0±2.5\*** |
| **n** | 6 | 6 | 2 | 6 | 2 | 6 | 6 | 6 |

**Table S2:**
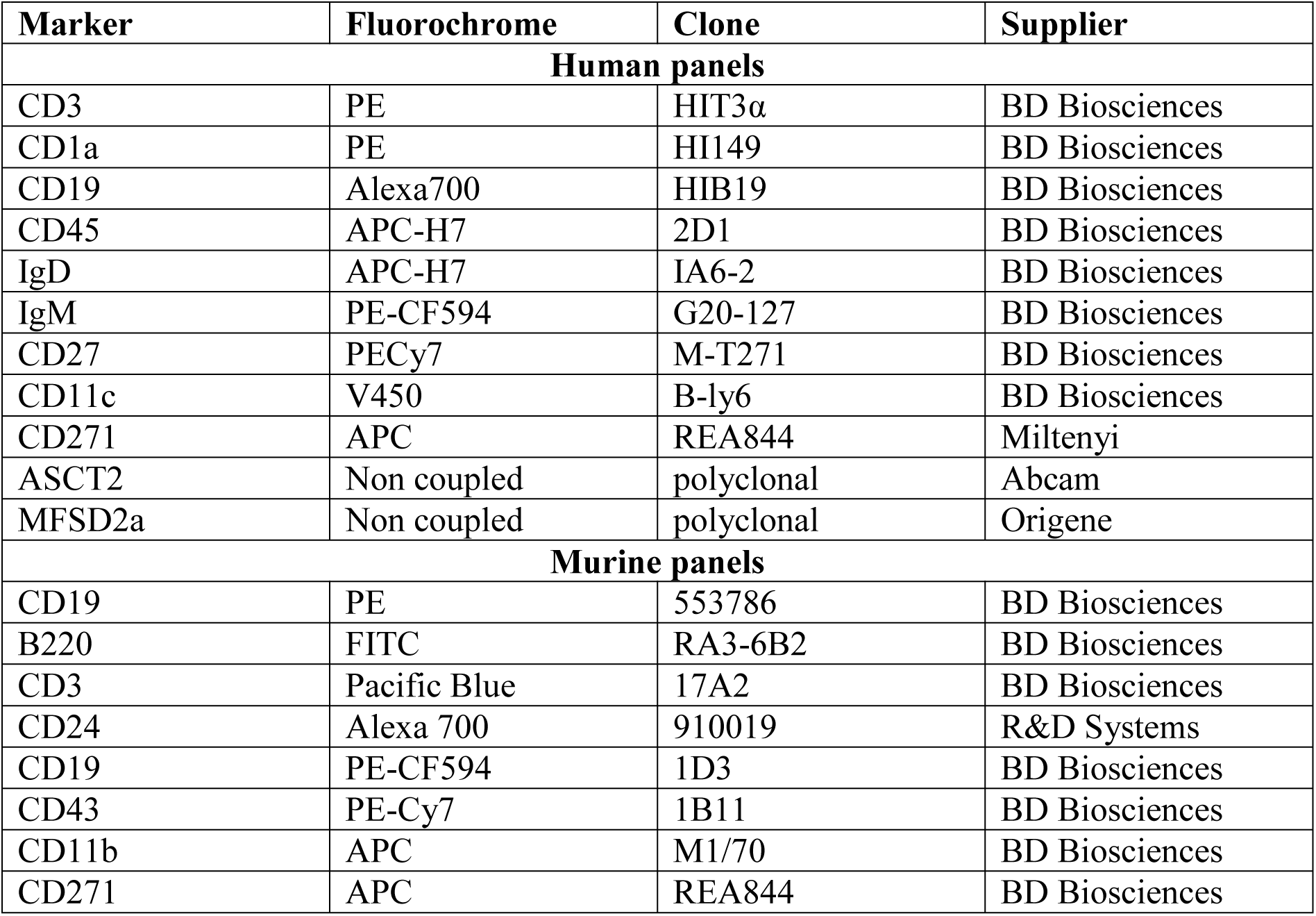
FACS antibody listing.

| **Marker** | **Fluorochrome** | **Clone** | **Supplier** |
| --- | --- | --- | --- |
| **Human panels** |  |  |  |
| CD3 | PE | HIT3α | BD Biosciences |
| CD1a | PE | HI149 | BD Biosciences |
| CD19 | Alexa700 | HIB19 | BD Biosciences |
| CD45 | APC-H7 | 2D1 | BD Biosciences |
| IgD | APC-H7 | IA6-2 | BD Biosciences |
| IgM | PE-CF594 | G20-127 | BD Biosciences |
| CD27 | PECy7 | M-T271 | BD Biosciences |
| CD11c | V450 | B-ly6 | BD Biosciences |
| CD271 | APC | REA844 | Miltenyi |
| ASCT2 | Non coupled | polyclonal | Abcam |
| MFSD2a | Non coupled | polyclonal | Origene |
| **Murine panels** |  |  |  |
| CD19 | PE | 553786 | BD Biosciences |
| B220 | FITC | RA3-6B2 | BD Biosciences |
| CD3 | Pacific Blue | 17A2 | BD Biosciences |
| CD24 | Alexa 700 | 910019 | R&D Systems |
| CD19 | PE-CF594 | 1D3 | BD Biosciences |
| CD43 | PE-Cy7 | 1B11 | BD Biosciences |
| CD11b | APC | M1/70 | BD Biosciences |
| CD271 | APC | REA844 | BD Biosciences |

**Table S3:**
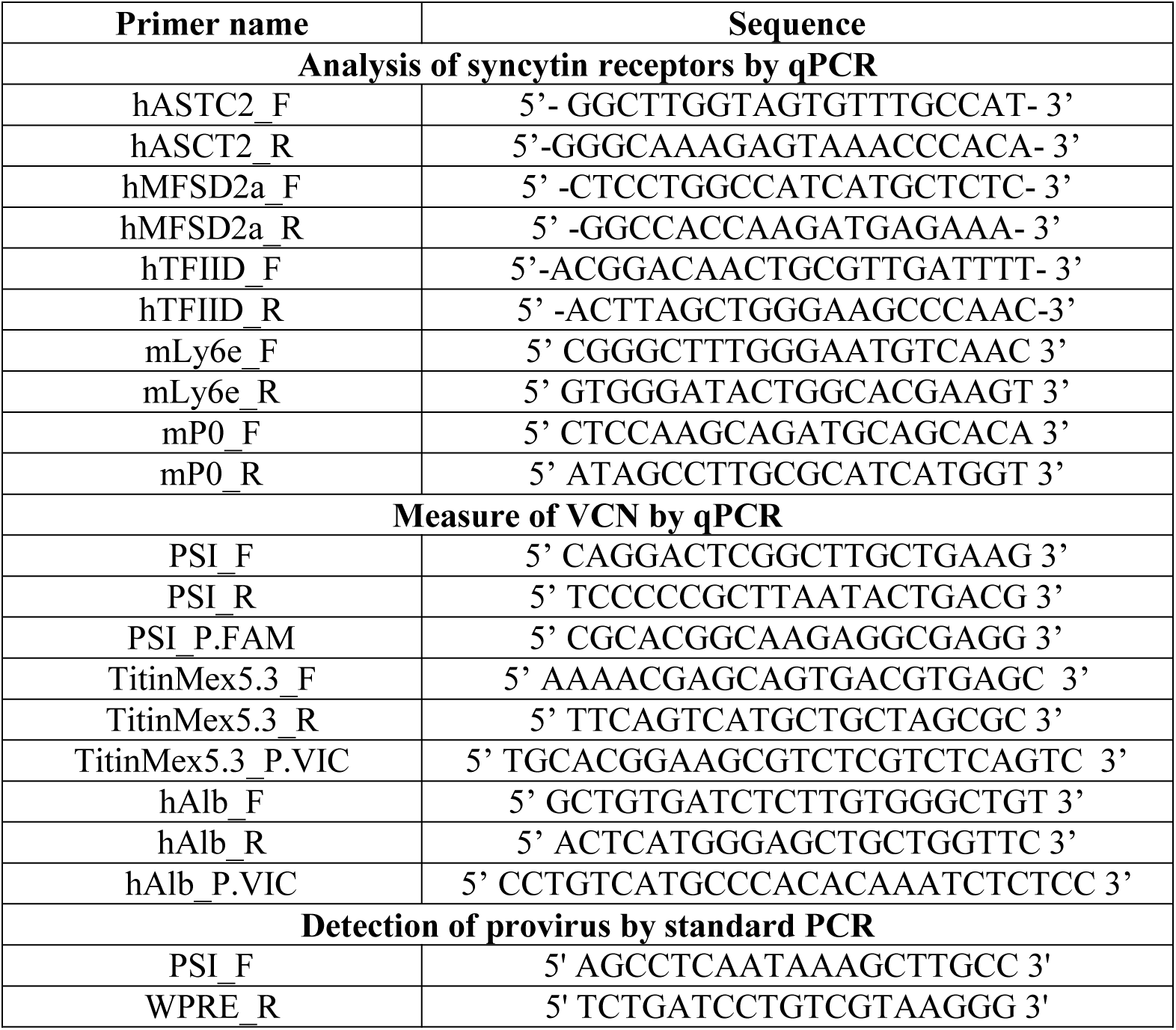
Primer sequences.

## Acknowledgments

We thank Dr. S. Fisson for providing A20.IIa cells and Dr. C. Poinsignon and S. Darocha for technical help. We are grateful to mothers donating cord blood and to Dr. Rigonnot and staff at the Centre Hospitalier Sud-Francilien for umbilical cord blood samples.

## Author contributions

YC and MF contributed equally planned and executed experiments and wrote the manuscript, KS and LM executed experiments, AG conceived and directed the study, wrote and edited the manuscript.

## Data and materials availability

Materials are available upon request and following signature of a MTA.

## References

1. Naldini L, Trono D, & Verma IM (2016) Lentiviral vectors, two decades later. Science 353(6304):1101–1102.

2. Galy A (2017) Major Advances in the Development of Vectors for Clinical Gene Therapy of Hematopoietic Stem Cells from European Groups over the Last 25 Years. Hum Gene Ther 28(11):964–971.

3. June CH & Sadelain M (2018) Chimeric Antigen Receptor Therapy. N Engl J Med 379(1):64–73.

4. Amirache F, et al. (2014) Mystery solved: VSV-G-LVs do not allow efficient gene transfer into unstimulated T cells, B cells, and HSCs because they lack the LDL receptor. Blood 123(9):1422–1424.

5. Cire S, et al. (2014) Immunization of mice with lentiviral vectors targeted to MHC class II+ cells is due to preferential transduction of dendritic cells in vivo. PLoS One 9(7):e101644.

6. Goyvaerts C, et al. (2017) Antigen-presenting cell-targeted lentiviral vectors do not support the development of productive T-cell effector responses: implications for in vivo targeted vaccine delivery. Gene Ther 24(6):370–375.

7. Milani M, et al. (2019) Phagocytosis-shielded lentiviral vectors improve liver gene therapy in nonhuman primates. Sci Transl Med 11(493).

8. Frank AM & Buchholz CJ (2019) Surface-Engineered Lentiviral Vectors for Selective Gene Transfer into Subtypes of Lymphocytes. Mol Ther Methods Clin Dev 12:19–31.

9. Dewannieux M & Heidmann T (2013) Endogenous retroviruses: acquisition, amplification and taming of genome invaders. Curr Opin Virol 3(6):646–656.

10. Dupressoir A, Lavialle C, & Heidmann T (2012) From ancestral infectious retroviruses to bona fide cellular genes: role of the captured syncytins in placentation. Placenta 33(9):663–671.

11. Dupressoir A, et al. (2005) Syncytin-A and syncytin-B, two fusogenic placenta-specific murine envelope genes of retroviral origin conserved in Muridae. Proc Natl Acad Sci U S A 102(3):725–730.

12. Dupressoir A, et al. (2009) Syncytin-A knockout mice demonstrate the critical role in placentation of a fusogenic, endogenous retrovirus-derived, envelope gene. Proc Natl Acad Sci U S A 106(29):12127–12132.

13. Lavialle C, et al. (2013) Paleovirology of ’syncytins’, retroviral env genes exapted for a role in placentation. Philos Trans R Soc Lond B Biol Sci 368(1626):20120507.

14. Cheynet V, et al. (2005) Synthesis, assembly, and processing of the Env ERVWE1/syncytin human endogenous retroviral envelope. J Virol 79(9):5585–5593.

15. Lavillette D, et al. (2002) The envelope glycoprotein of human endogenous retrovirus type W uses a divergent family of amino acid transporters/cell surface receptors. J Virol 76(13):6442–6452.

16. Blond JL, et al. (2000) An envelope glycoprotein of the human endogenous retrovirus HERV-W is expressed in the human placenta and fuses cells expressing the type D mammalian retrovirus receptor. J Virol 74(7):3321–3329.

17. Esnault C, et al. (2008) A placenta-specific receptor for the fusogenic, endogenous retrovirus-derived, human syncytin-2. Proc Natl Acad Sci U S A 105(45):17532–17537.

18. Dupressoir A, et al. (2011) A pair of co-opted retroviral envelope syncytin genes is required for formation of the two-layered murine placental syncytiotrophoblast. Proc Natl Acad Sci U S A 108(46):E1164–1173.

19. Bacquin A, et al. (2017) A Cell Fusion-Based Screening Method Identifies Glycosylphosphatidylinositol-Anchored Protein Ly6e as the Receptor for Mouse Endogenous Retroviral Envelope Syncytin-A. J Virol 91(18).

20. Frese S, et al. (2015) Long-Term Endurance Exercise in Humans Stimulates Cell Fusion of Myoblasts along with Fusogenic Endogenous Retroviral Genes In Vivo. PLoS One 10(7):e0132099.

21. Redelsperger F, et al. (2016) Genetic Evidence That Captured Retroviral Envelope syncytins Contribute to Myoblast Fusion and Muscle Sexual Dimorphism in Mice. PLoS Genet 12(9):e1006289.

22. Coudert AE, et al. (2019) Role of the captured retroviral envelope syncytin-B gene in the fusion of osteoclast and giant cell precursors and in bone resorption, analyzed ex vivo and in vivo in syncytin-B knockout mice. Bone Rep 11:100214.

23. Mangeney M, et al. (2007) Placental syncytins: Genetic disjunction between the fusogenic and immunosuppressive activity of retroviral envelope proteins. Proc Natl Acad Sci U S A 104(51):20534–20539.

24. An DS, Xie Y, & Chen IS (2001) Envelope gene of the human endogenous retrovirus HERV-W encodes a functional retrovirus envelope. J Virol 75(7):3488–3489.

25. Blaise S, Ruggieri A, Dewannieux M, Cosset FL, & Heidmann T (2004) Identification of an envelope protein from the FRD family of human endogenous retroviruses (HERV-FRD) conferring infectivity and functional conservation among simians. J Virol 78(2):1050–1054.

26. Fenard D, et al. (2013) Vectofusin-1, a new viral entry enhancer, strongly promotes lentiviral transduction of human hematopoietic stem cells. Mol Ther Nucleic Acids 2:e90.

27. Majdoul S, et al. (2016) Molecular Determinants of Vectofusin-1 and Its Derivatives for the Enhancement of Lentivirally Mediated Gene Transfer into Hematopoietic Stem/Progenitor Cells. J Biol Chem 291(5):2161–2169.

28. Fenard D, et al. (2013) Infectivity enhancement of different HIV-1-based lentiviral pseudotypes in presence of the cationic amphipathic peptide LAH4-L1. J Virol Methods 189(2):375–378.

29. Charrier S, et al. (2011) Quantification of lentiviral vector copy numbers in individual hematopoietic colony-forming cells shows vector dose-dependent effects on the frequency and level of transduction. Gene Ther 18(5):479–487.

30. Kaminski DA, Wei C, Qian Y, Rosenberg AF, & Sanz I (2012) Advances in human B cell phenotypic profiling. Front Immunol 3:302.

31. Recher M, et al. (2011) IL-21 is the primary common gamma chain-binding cytokine required for human B-cell differentiation in vivo. Blood 118(26):6824–6835.

32. Marin M, Tailor CS, Nouri A, & Kabat D (2000) Sodium-dependent neutral amino acid transporter type 1 is an auxiliary receptor for baboon endogenous retrovirus. J Virol 74(17):8085–8093.

33. Oburoglu L, et al. (2014) Glucose and glutamine metabolism regulate human hematopoietic stem cell lineage specification. Cell Stem Cell 15(2):169–184.

34. Girard-Gagnepain A, et al. (2014) Baboon envelope pseudotyped LVs outperform VSV-G-LVs for gene transfer into early-cytokine-stimulated and resting HSCs. Blood 124(8):1221–1231.

35. Piovan C, et al. (2017) Vectofusin-1 Promotes RD114-TR-Pseudotyped Lentiviral Vector Transduction of Human HSPCs and T Lymphocytes. Mol Ther Methods Clin Dev 5:22–30.

36. Radek C, et al. (2019) Vectofusin-1 improves transduction of primary human cells with diverse retroviral and lentiviral pseudotypes, enabling robust, automated closed-system manufacturing. Hum Gene Ther.

37. Hummel J, Kammerer U, Muller N, Avota E, & Schneider-Schaulies S (2015) Human endogenous retrovirus envelope proteins target dendritic cells to suppress T-cell activation. Eur J Immunol 45(6):1748–1759.

38. Lokossou AG, et al. (2019) Endogenous retrovirus-encoded Syncytin-2 contributes to exosome-mediated immunosuppression of T cells. Biol Reprod.

39. Levy C, et al. (2016) Baboon envelope pseudotyped lentiviral vectors efficiently transduce human B cells and allow active factor IX B cell secretion in vivo in NOD/SCIDgammac(-/-) mice. J Thromb Haemost 14(12):2478–2492.

40. Bernadin O, et al. (2019) Baboon envelope LVs efficiently transduced human adult, fetal, and progenitor T cells and corrected SCID-X1 T-cell deficiency. Blood Adv 3(3):461–475.

41. Frecha C, et al. (2009) Efficient and stable transduction of resting B lymphocytes and primary chronic lymphocyte leukemia cells using measles virus gp displaying lentiviral vectors. Blood 114(15):3173–3180.

42. Funke S, et al. (2008) Targeted cell entry of lentiviral vectors. Mol Ther 16(8):1427–1436.

43. Kneissl S, et al. (2013) CD19 and CD20 targeted vectors induce minimal activation of resting B lymphocytes. PLoS One 8(11):e79047.

44. Nguyen LN, et al. (2014) Mfsd2a is a transporter for the essential omega-3 fatty acid docosahexaenoic acid. Nature 509(7501):503–506.

45. Kichler A, Mason AJ, Marquette A, & Bechinger B (2013) Histidine-rich cationic amphipathic peptides for plasmid DNA and siRNA delivery. Methods Mol Biol 948:85–103.

46. Ingrao D, Majdoul S, Seye AK, Galy A, & Fenard D (2014) Concurrent measures of fusion and transduction efficiency of primary CD34+ cells with human immunodeficiency virus 1-based lentiviral vectors reveal different effects of transduction enhancers. Hum Gene Ther Methods 25(1):48–56.

47. Vermeer LS, et al. (2017) Vectofusin-1, a potent peptidic enhancer of viral gene transfer forms pH-dependent alpha-helical nanofibrils, concentrating viral particles. Acta Biomater 64:259–268.

48. Kutner RH, Zhang XY, & Reiser J (2009) Production, concentration and titration of pseudotyped HIV-1-based lentiviral vectors. Nat Protoc 4(4):495–505.

